# HuR inhibition attenuates hypertension and fibrosis in chronic kidney disease

**DOI:** 10.64898/2026.02.23.707592

**Authors:** Lili Zhuang, Zhou Wang, Ziwei Fu, Sohom Mookherjee M. Tech, J. David Symons, Jeffrey Aube, Xiaoqing Wu, Liang Xu, Yufeng Huang

## Abstract

**Background:** Elevated RNA-binding protein HuR has been reported in patients with chronic kidney disease (CKD), but its specific pathogenic role remains unclear. Here, we investigated HuR involvement in progressive CKD induced by deoxycorticosterone acetate (DOCA) plus angiotensin II (Ang II) in mice and evaluated the therapeutic efficacy and mechanisms of the HuR inhibitor KH3.

**Methods:** Adult male mice underwent uninephrectomy and were subjected to DOCA + Ang II infusion with 1% NaCl in drinking water. Mice were then treated with KH3 or vehicle for 3 weeks. Control mice received saline injections without DOCA and Ang II infusion.

**Results:** DOCA + Ang II infusion markedly increased HuR expression in circulating exosomes and kidney tissues, which was attenuated by KH3. KH3 halted the progression of albuminuria and improved renal function, and reduced kidney hypertrophy and glomerular and tubulointerstitial fibrosis compared with untreated DOCA + Ang II mice. These improvements were associated with reduced podocyte and tubular injury. KH3 also decreased renal macrophage infiltration and suppressed NF-κBp65, Nox2, AKT phosphorylation, TGF-β1, and Wisp1, consistent with reduced inflammation, oxidative stress, and fibrosis. In addition, KH3 partially lowered arterial blood pressure in DOCA + Ang II–infused mice, an effect that may involve suppression of SGLT2-associated profibrotic vascular responses, as supported by studies in cultured VSMCs and mesenteric resistance arteries.

**Conclusions:** HuR is upregulated in DOCA + Ang II–induced renal and vascular injury and contributes to hypertensive, inflammatory, oxidative, and fibrotic responses in CKD. Pharmacologic inhibition of HuR-RNA interactions represents a promising therapeutic strategy for CKD.

**NOVELTY AND RELEVANCE:** *What Is New?:* This study identifies the RNA-binding protein HuR (ELAVL1) as a previously unrecognized upstream post-transcriptional regulator of blood pressure in hypertensive chronic kidney disease. We demonstrate for the first time that pharmacologic disruption of HuR–RNA interactions lowers arterial blood pressure in vivo. In addition, we uncover a novel HuR–SGLT2–vascular smooth muscle cell (VSMC) signaling axis, revealing that HuR regulates inducible vascular SGLT2 expression and Ang II–mediated vasoconstrictive responses.

*What Is Relevant?:* Hypertension in CKD arises from integrated renal and vascular dysfunction that is incompletely controlled by current therapies. Our findings are highly relevant because we identify HuR as a nodal post-transcriptional regulator that coordinates renal injury, vascular inflammation, and smooth muscle contractility, rather than acting within a single cell type or signaling pathway.

*Clinical and Pathophysiological Implication:* These data support a model in which HuR-driven RNA regulatory programs amplify Ang II–dependent vascular hypercontractility and hypertension in CKD. Therapeutic targeting of HuR–RNA interactions represents a novel antihypertensive strategy that may complement renin–angiotensin–aldosterone system (RAAS) blockade and provides mechanistic insight into the blood pressure–lowering and vascular protective effects of SGLT2 inhibitors, including in non-diabetic CKD.

## INTRODUCTION

Chronic kidney disease (CKD) represents a major global public health burden, with steadily increasing prevalence worldwide ^1^. Although current therapies—including renin–angiotensin system inhibitors (ACE inhibitors/ARBs), nonsteroidal mineralocorticoid receptor antagonists (e.g. finerenone), sodium–glucose cotransporter 2 (SGLT2) inhibitors—can slow disease progression, they do not halt it, and many patients ultimately progress to end-stage kidney disease (ESKD). The upstream molecular regulators that integrate renal injury, vascular dysfunction, and blood pressure control in CKD remain incompletely defined. Thus, there remains a critical unmet need for mechanism-based therapies that target upstream drivers of CKD progression.

Human antigen R (HuR; ELAVL1) is an RNA-binding protein that post-transcriptionally regulates the stability and translation of numerous inflammatory, oxidative stress–related, and fibrotic mediators by binding adenylate–uridylate–rich elements (AREs) within 3′ untranslated regions (3′UTRs) of target mRNAs. Aberrant HuR upregulation has been documented in a wide range of inflammatory and fibrotic disorders, including kidney diseases ^2–12^. Analysis of human transcriptomic datasets (Nephroseq) further supports elevated renal HuR mRNA expression in patients with CKD.

Consistent with these observations, our prior studies demonstrated marked HuR upregulation in experimental models of chronic glomerulonephritis, ischemia-induced tubular fibrosis, and type 2 diabetic nephropathy ^6, 13, 14^. We also developed a potent small-molecule HuR inhibitor, KH3, that disrupts HuR–ARE interactions ^15, 16^ and significantly attenuates renal inflammation, oxidative stress, and tissue injury in these models.

In the present study, we investigated the pathogenic role of HuR in progressive hypertensive CKD induced by DOCA + Ang II infusion in mice. We further evaluated the therapeutic efficacy and mechanistic actions of HuR inhibition using KH3, with the goal of defining HuR as a tractable upstream regulator and therapeutic target in CKD progression.

## MATERIALS AND METHODS

### Data availability

The data that support the findings of this study are available from the corresponding author upon reasonable request.

### Reagents

The HuR inhibitor KH3 was synthesized as previously described ^15, 16^. For in vivo studies, KH3 powder was dissolved in PBS containing 5% ethanol and 5% Tween-80. For in vitro assays, KH3 was prepared as a 20 mM DMSO stock and diluted in PBS as needed ^16^. Dapagliflozin was kindly provided by AstraZeneca (Mölndal, Sweden) as described previously ^17^ and dissolved in PBS for in vitro assays. Unless otherwise stated, all other reagents were purchased from Sigma-Aldrich (St. Louis, MO, USA).

### Study 1. In vivo changes and inhibition of HuR in a DOCA + Ang II–induced CKD model

#### Animals

Animal housing and care followed the NIH Guide for the Care and Use of Laboratory Animals. All procedures were approved by the University of Utah Institutional Animal Care and Use Committee.

#### Experimental design

Twenty-six male C57BL/6 mice (Jackson Laboratory, Bar Harbor, ME, USA) underwent uninephrectomy under isoflurane anesthesia at 10 weeks of age. Two weeks later, mice were randomized into three groups: (i) Normal control (n = 8): no infusion and standard drinking water. (ii) DOCA + angiotensin II (Ang II) (n = 9): subcutaneous implantation of a 25 mg DOCA pellet (Innovative Research, FL, USA) plus Ang II infusion (1 ng/min/g body weight) via osmotic mini pump (Alzet 1004, Durect, Cupertino, CA, USA), with 1% NaCl in drinking water for 3 weeks. (iii) DOCA + Ang II + KH3 (n = 9): same as group ii, plus KH3 (40 mg/kg/day, i.p.), initiated 3 days after starting DOCA + Ang II infusion. Model induction was performed as previously reported ^18^.

Systolic blood pressure was measured at baseline and before sacrifice in conscious, trained mice (≈10 AM) using the tail-cuff method (MC4000 multi-channel system, Hatteras Instruments, Cary, NC, USA). Body weight, 24-hour water intake, and urine output were recorded weekly using metabolic cages beginning before treatment. Urine albumin excretion (UAE) and the albumin-to-creatinine ration (ACR) were measured using the DCA 2000+ microalbumin/creatinine kit (Bayer HealthCare, Elkhart, IN, USA).

Mice were euthanized under isoflurane anesthesia. Blood was collected via cardiac puncture for plasma BUN/creatinine (Cr) measurement and exosome isolation. Kidneys were perfused via the heart with 30 mL cold sterile PBS, weighed, and processed for histology and molecular analyses. Cortical tissue was snap-frozen in 2-methylbutane at −80°C or fixed in 10% neutral buffered formalin. Additional cortex was stored for Western blotting or processed in TRIzol (Invitrogen, Carlsbad, CA, USA) for RNA isolation. Thoracic and abdominal aortas were isolated and processed in TRIzol for RNA isolation.

#### Measurement of renal function

Plasma blood urea nitrogen (BUN) and creatinine (Cr) were measured using the QuantiChrom urea assay kit (DIUR-100; BioAssay Systems, Hayward, CA, USA) and the creatinine assay kit (80350; Crystal Chem Inc., Elk Grove Village, IL, USA), respectively.

#### Plasma exosome isolation and circulating HuR measurement

Exosomes were isolated using the ExoQuick Ultra extracellular vehicle (EV) kit (EQYKTRA-20A-1, System Biosciences, Palo Alto, CA, USA) per manufacturer instructions. Briefly, 0.25 mL plasma per mouse was processed. Protein concentration was measured by the bicinchoninic acid (BCA) protein assay (Pierce Biotechnology, Rockford, IL, USA). Equal exosomal protein (20 µg) was separated by SDS-PAGE (4–12% gradient gel) (Invitrogen) and immunoblotted for HuR and CD63.

#### Histological analysis

Paraffin-embedded kidney sections (4 µm) were stained with periodic acid-Schiff (PAS) and Masson’s trichrome (TRI). PAS-positive mesangial matrix was quantified in a blinded manner using a computer-assisted method as previously described ^18^. Trichrome-stained collagen deposition was quantified in ten non-overlapping cortical fields per section at 200x using ImageJ and expressed as a percentage of the total tissue area.

#### Immunofluorescence and immunohistochemistry

HuR immunofluorescence was performed on paraffin sections as previously described ^6, 13^. Alexa Fluor 488–conjugated wheat germ agglutinin (WGA) was used to delineate glomeruli and tubules; nuclei were counterstained with DAPI. Control slides without primary antibody showed no staining.

Nephrin and podocin staining was performed on frozen sections. α-SMA, fibronectin (FN), collagen III (Col-III), and SGLT2 staining was performed on paraffin sections as previously described ^19–21^. Macrophages were detected using rat anti-mouse F4/80 and Cy3-conjugated secondary antibody. F4/80+ cells were quantified in five randomly selected cortical fields per section at 200x. Antibody details are provided in Supplemental Table S1.

#### Western blot analysis

Renal cortex (20 mg) was homogenized in lysis buffer (Cell Signaling Technology, Inc. Danvers, MA, USA) containing 1% NP-40, 1 mM PMSF, and protease inhibitors. Protein (30 µg) was resolved on 4–12% SDS-PAGE gels (Invitrogen) and probed for HuR, FN, α-SMA, vimentin, PDGFR-ß, NF-κBp65, Nox2, phospho-AKT, Kim-1, OPN, TGFß1, Wisp1, SGLT2, GLP-1R, ENaC subunits (α/β/γ), GAPDH, and β-actin (antibodies as listed in Supplemental Table S1). Bands were detected by ECL and quantified by ImageJ. Protein levels were normalized to β-actin or GAPDH and expressed as fold-change versus controls. Blots were performed at least three times.

#### RNA isolation and RT-qPCR

Total RNA was extracted using TRIzol, reverse-transcribed (SuperScript III (Invitrogen)) and quantified by SYBR Green (Invitrogen) RT-qPCR on ABI 7900 as previously described ^13, 19^. Target gene expression was normalized to GAPDH. Primer sequences are listed in Supplemental Table S2. Product specificity was confirmed by agarose gel electrophoresis.

### Study 2. HuR inhibition and SGLT2 regulation in primary mouse VSMCs

Primary mouse proximal tubular cells (mPTCs) and arterial VSMCs (mVSMCs) were prepared and validated as previously described ^22, 23^. mPTCs served as positive controls for SGLT2 expression. Cells (≤ passage 10) were maintained in DMEM + 10% FBS with penicillin/streptomycin at 37°C and 5% CO₂. Sub-confluent cells were serum-starved for 24 hours before stimulation. Ang II (10⁻⁷ M), TGF-β1 (5 ng/mL), KH3 (10 µM), and dapagliflozin (Dapa, 20 µM) were used as indicated. Experiments were performed in duplicate and repeated three times.

### Study 3. Mesenteric resistance artery reactivity

Second-order mesenteric resistance arteries were studied ex vivo using isometric tension techniques (PMID: 33890822). Four arterial segments from each mouse were examined per day on four separate days. After mounting the arteries and determining Lmax (Supplementary data Table S3), contractile responses to potassium chloride (KCl) and phenylephrine (PE) were assessed. Arteries were then incubated for 2 h with vehicle (PBS), KH3 (10 µM), dapagliflozin (Dapa, 20 µM), or KH3 + dapagliflozin. Responses to Ang II were then evaluated, followed by assessing endothelium-dependent vasorelaxation to acetylcholine (Ach) and endothelium-independent vasorelaxation to sodium nitroprusside (SNP). Details of the experimental protocol are provided in the Supplemental Methods.

### Statistical analyses

All non-vascular data are presented as mean ± SD. Sample size calculations were performed using Power Analysis software (www.clincalc.com) based on urinary albumin excretion from a pilot study (2310 ± 1445 µg/day, n = 4), which was used as the primary outcome measure. Power calculations assumed a two-sided, two-sample t test with equal variance, a detectable effect size of 2.2 standard deviations (d = 2.2), α = 0.05, and a target power of 95% (0.95). The estimated sample size was 6 mice per group. To account for potential attrition, 8 or more mice per group were included. Group comparisons were performed using two-way ANOVA followed by Student–Newman–Keuls or Dunnett post hoc tests. P < 0.05 was considered significant.

Vascular reactivity data are presented as mean ± SEM. Normality was assessed in GraphPad Prism 9. Comparisons among groups across doses were analyzed by two-way repeated-measures ANOVA with Tukey multiple comparisons as appropriate. Statistical details are provided in each figure legend.

## RESULTS

### Elevated HuR in DOCA + Ang II–induced CKD

As shown in Fig. 1 A–B, HuR was detectable in circulating exosomes, and its levels were significantly increased in mice with DOCA + Ang II–induced hypertensive CKD. In kidneys, HuR immunofluorescence intensity was markedly increased, with prominent nuclear localization and additional cytoplasmic staining in glomerular, tubular, and vascular cells (Fig. 1C, arrows). HuR-positive staining was also evident in tubulointerstitial infiltrating cells (Fig. 1C, arrows). In contrast, control kidneys showed predominantly nuclear HuR with weak glomerular and tubular staining. Consistent with these findings, immunoblotting demonstrated increased total renal HuR protein in diseased kidneys (Fig. 1 D–E). KH3 treatment markedly reduced HuR levels in both circulating exosomes and kidney tissue toward control levels (Fig. 1 A–E). These data indicate robust systemic and renal HuR upregulation in hypertensive CKD and effective suppression of disease-associated HuR overexpression by KH3.

**Figure 1.**
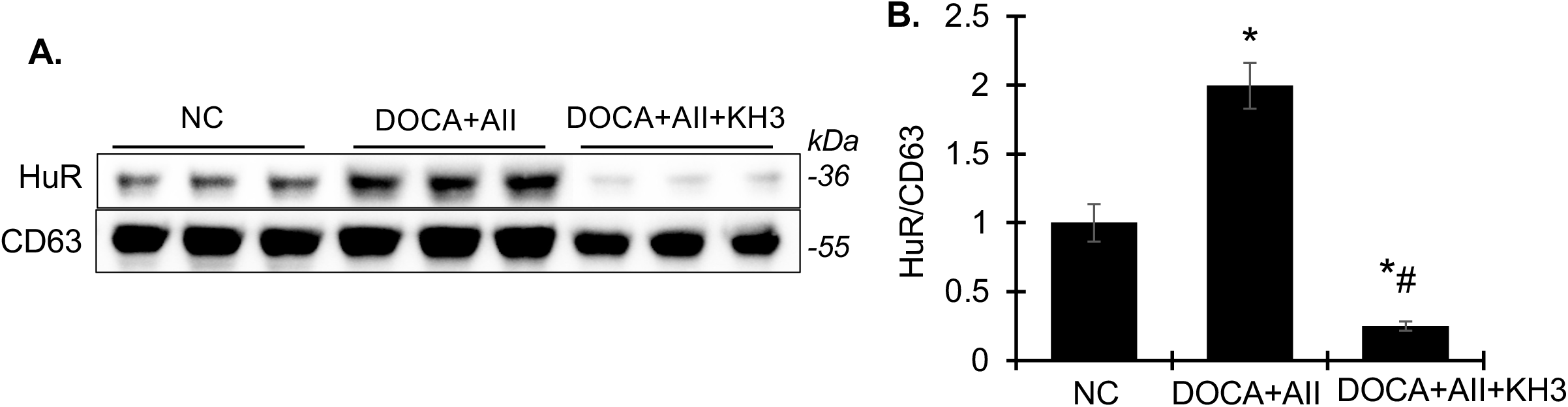
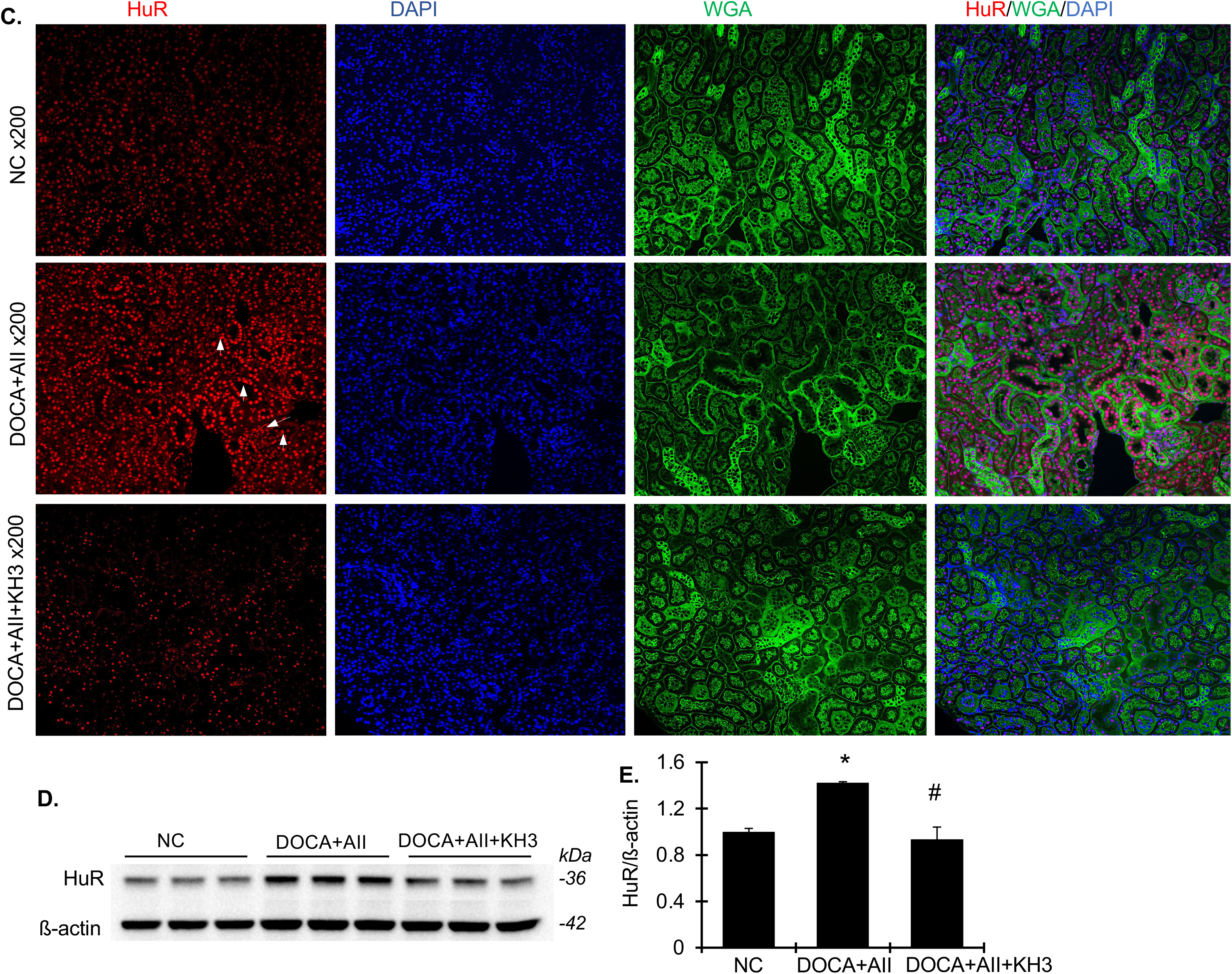
Treatment with KH3 reversed elevated HuR expression in circulating exosomes and kidney tissue in DOCA + Ang II-induced CKD mice. **(A)** Representative Western blots showing HuR and the exosomal marker CD63 in circulating plasma exosomes isolated from normal control (NC), DOCA + Ang II–treated (DOCA + AII), and KH3-treated DOCA + Ang II mice (DOCA + AII + KH3). **(B)** Densitometric quantification of exosomal HuR protein levels normalized to CD63. **(C)** Representative immunofluorescence images of renal cortex sections stained for HuR (red), nuclei (DAPI, blue), and wheat germ agglutinin (WGA, green) at ×200 magnification. Compared with NC kidneys, DOCA + Ang II–injured kidneys exhibited increased HuR expression with prominent cytoplasmic localization in glomerular, tubular, and vascular cells (arrows), which was markedly reduced by KH3 treatment. **(D)** Representative Western blots showing HuR and β-actin expression in whole kidney lysates. **(E)** Densitometric quantification of renal HuR protein levels normalized to β-actin. Protein expression levels are expressed relative to NC, which was set to unity. KH3 treatment significantly reduced circulating exosomal HuR levels as well as renal glomerular and tubular HuR expression in DOCA + Ang II–injured mice. Data are presented as mean ± SD. *P<0.05, vs. NC; #P<0.05, vs. DOCA+AII.

### KH3 attenuates DOCA + Ang II–induced renal disease

#### Systemic parameters

Baseline and terminal parameters are summarized in Fig. 2. All mice survived to end of study. Over 3 weeks, control mice gained ∼1.08 g, whereas DOCA + Ang II mice lost ∼0.78 g; body weight remained stable in KH3-treated mice (Fig. 2A). DOCA + Ang II induced marked hypertension, as assessed by tail-cuff plethysmograph, which was partially but significantly attenuated by KH3 (Fig. 2B). Water intake and urine output increased beginning in week 1 and persisted through week 3; both were significantly reduced by KH3 (Fig. 2 C–D). DOCA + Ang II infusion also caused progressive albuminuria determined by urinary albumin-to-creatinine ratio (ACR) and 24-hour urinary albumin excretion (UAE); KH3 significantly attenuated albuminuria (Fig. 2 E–F). Plasma BUN and Cr levels were elevated in hypertensive mice and markedly improved by KH3 (Fig. 2 G–H).

**Figure 2.**
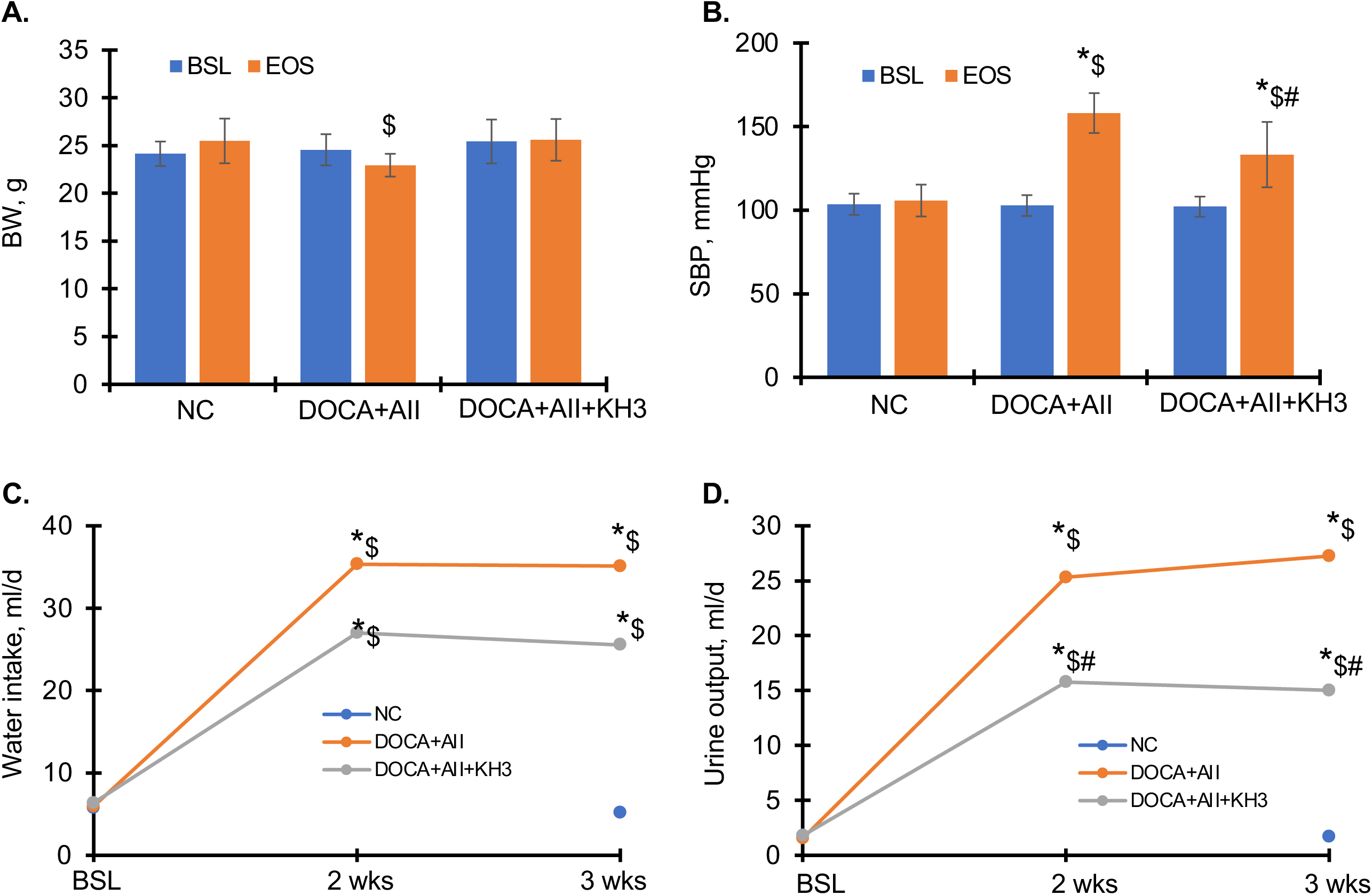
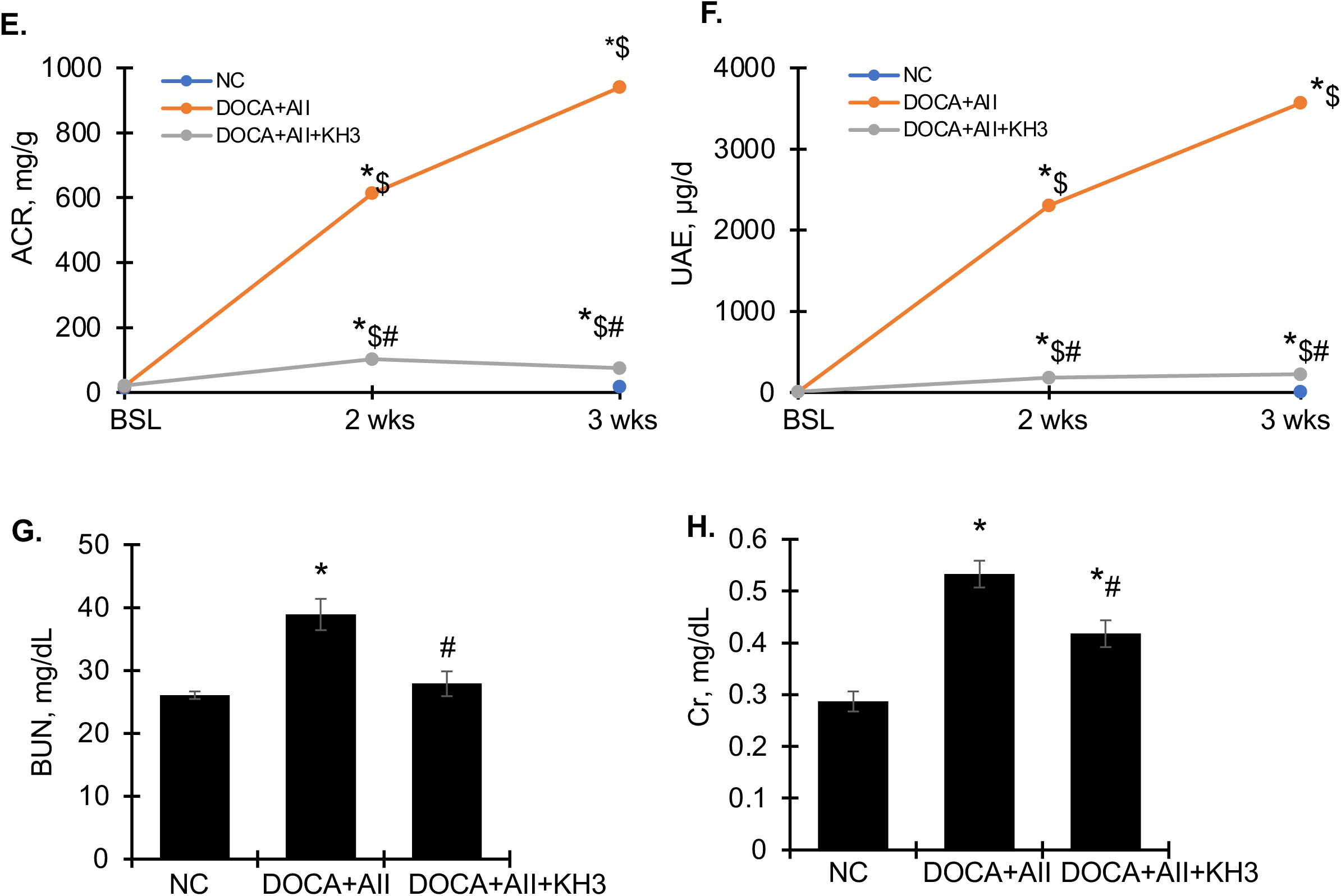
Effects of KH3 on blood pressure and renal function in DOCA + Ang II-induced CKD mice. (**A-H**) Body weight (**A**), systolic blood pressure (**B**), daily water intake (**C**), urine output (**D**), daily urinary albumin excretion (UAE) (**E**), and urinary albumin-to-creatinine ratio (ACR) (**F**) and plasma BUN (**G**) and Cr levels (**H**) in in normal control (NC), DOCA + Ang II–treated (DOCA + AII), and KH3-treated DOCA + Ang II mice. Data are presented as mean ± SD. *P<0.05, vs. NC; #P<0.05, vs. DOCA+AII.

#### HuR inhibition improves renal histology and fibrosis

After 3 weeks, kidney weight increased in DOCA + Ang II mice and was modestly reduced by KH3 (Fig. 3A). PAS staining demonstrated glomerular extracellular matrix (ECM) expansion in DOCA + Ang II-infused kidneys (Fig. 3B), confirmed by quantitative analysis (Fig. 3D). DOCA + Ang II-infused kidneys also showed tubular dilation, casts, interstitial mononuclear infiltration, and increased collagen deposition (Fig. 3 C, E–G). These abnormalities were markedly attenuated by KH3.

**Figure 3.**
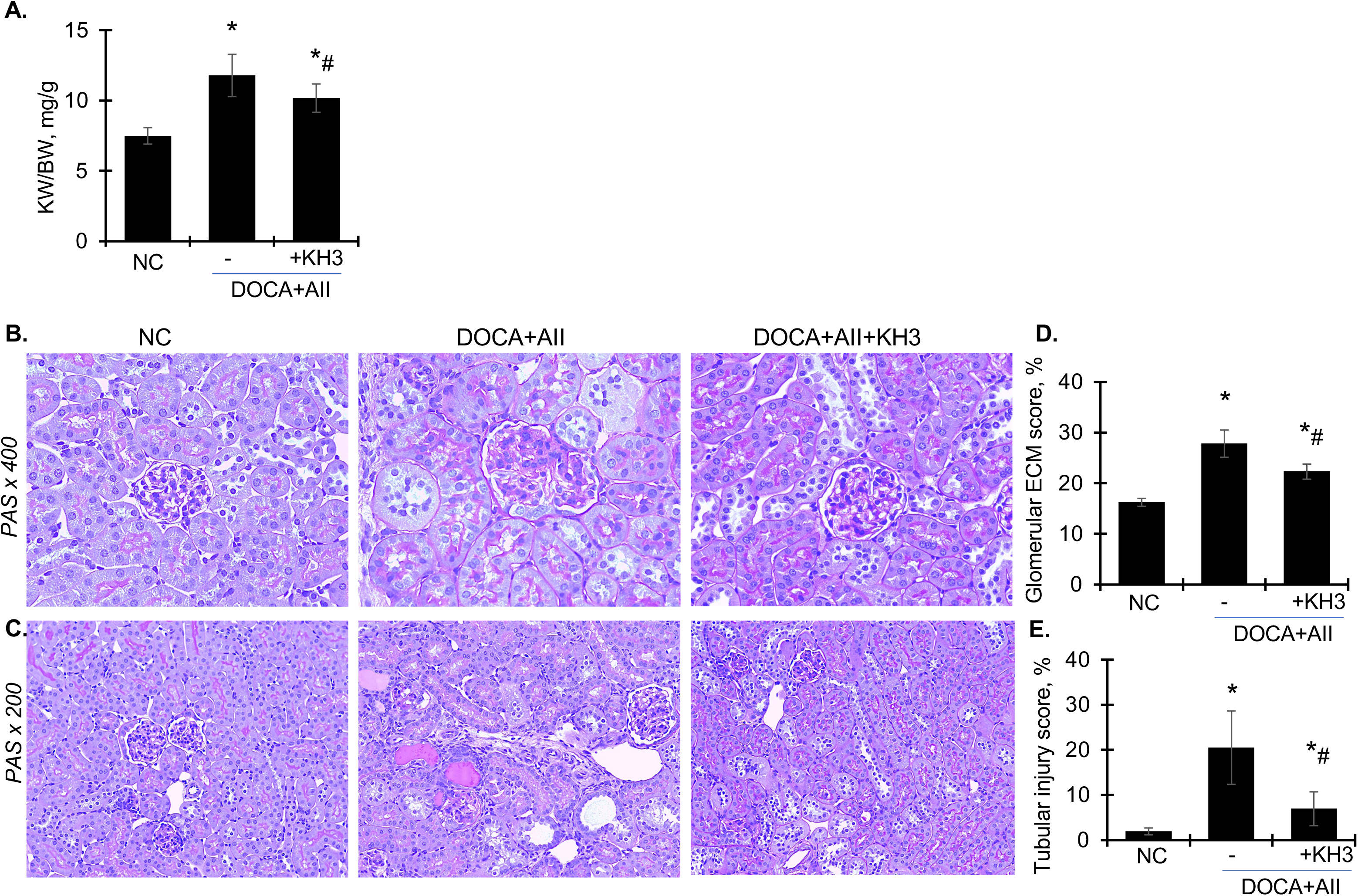
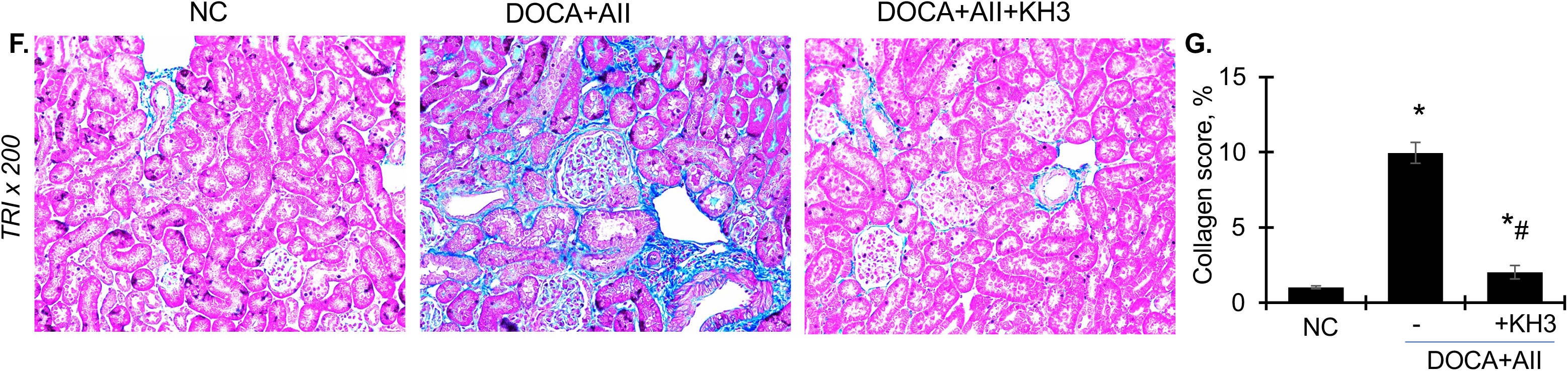
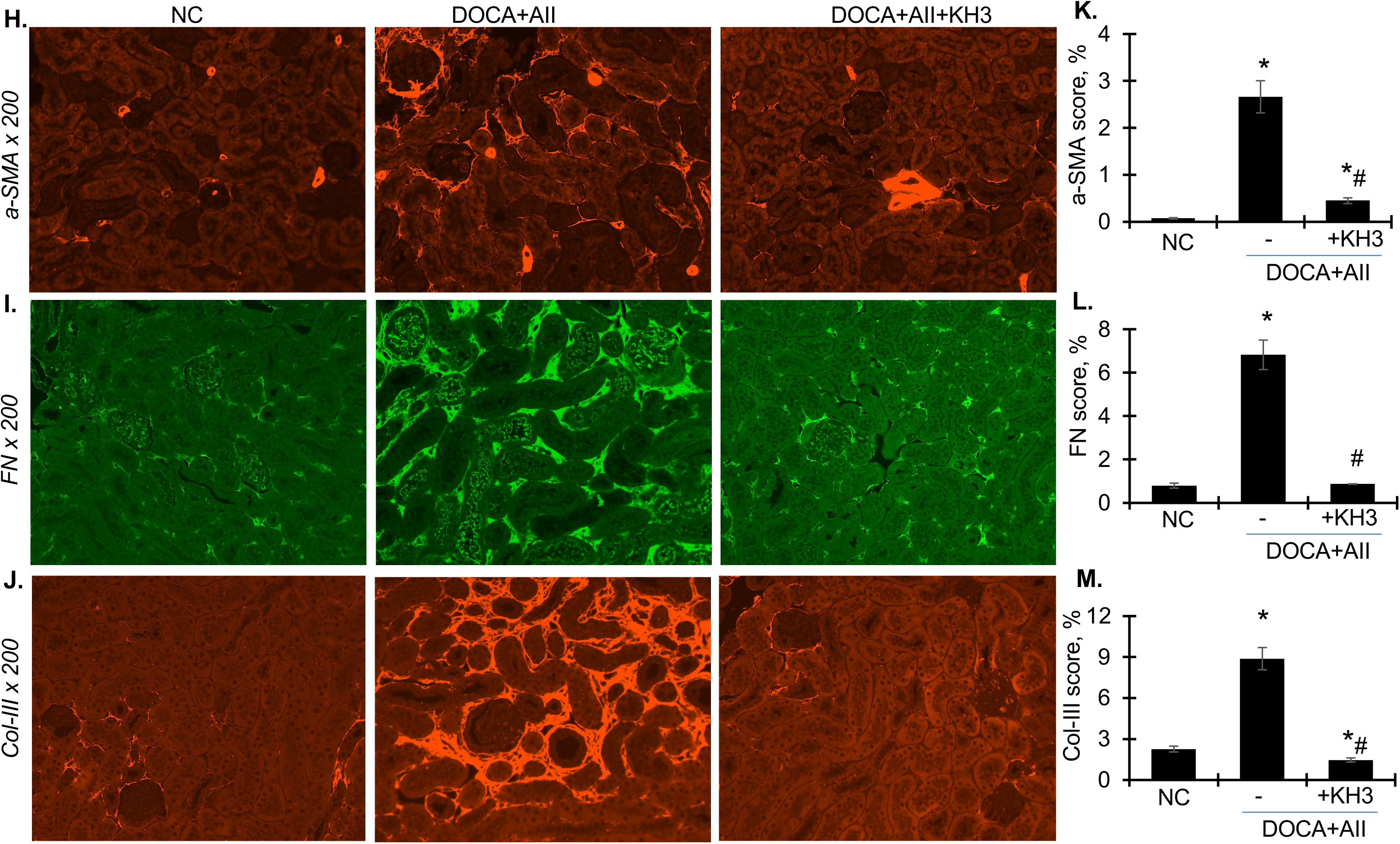
Treatment with KH3 ameliorates kidney hypertrophy, glomerular and tubular injury, and fibrosis in DOCA + Ang II-induced CKD mice. (**A-G**) Kidney weight (**A**) and representative histological images of kidney sections stained with PAS staining to assess glomerular and tubular injury (**B**, x400; **C**, x200) and Masson’s Trichrome staining to evaluate collagen deposition (**F**, x200). Quantitative analyses of glomerular extracellular matrix (ECM) accumulation (**D**), tubular injury score (**E**), and interstitial collagen deposition (**G**) were performed using image-J. (**H-M**) Representative confocal images of kidney sections immunostained for a-SMA (red) (**H**), FN (green) (**I**), and collagen III (Col-III, red) (**J**). Images were acquired at x200 magnification. Quantification of a-SMA (**K**), FN (**L**), and Col-III (**M**) staining was performed using image-J. Data are presented as mean ± SD. *P<0.05 vs normal control (NC). #P<0.05, vs. DOCA+AII.

Immunostaining showed increased interstitial α-SMA beyond vascular structures (indicating myofibroblast activation; Fig. 3 H, K), increased FN deposition in glomerular and tubulointerstitial compartments (Fig. 3 I, L), and increased Col-III along tubular basement membranes and interstitium (Fig. 3 J, M). KH3 markedly reduced these fibrotic changes (Fig. 3 B–M). Consistently, KH3 reduced renal protein levels of FN, α-SMA, vimentin, and PDGFR-β (Fig. 4 A–E) and decreased mRNA levels of FN, α-SMA, Col-I, and Col-III (Fig. 4 F–I).

**Figure 4.**
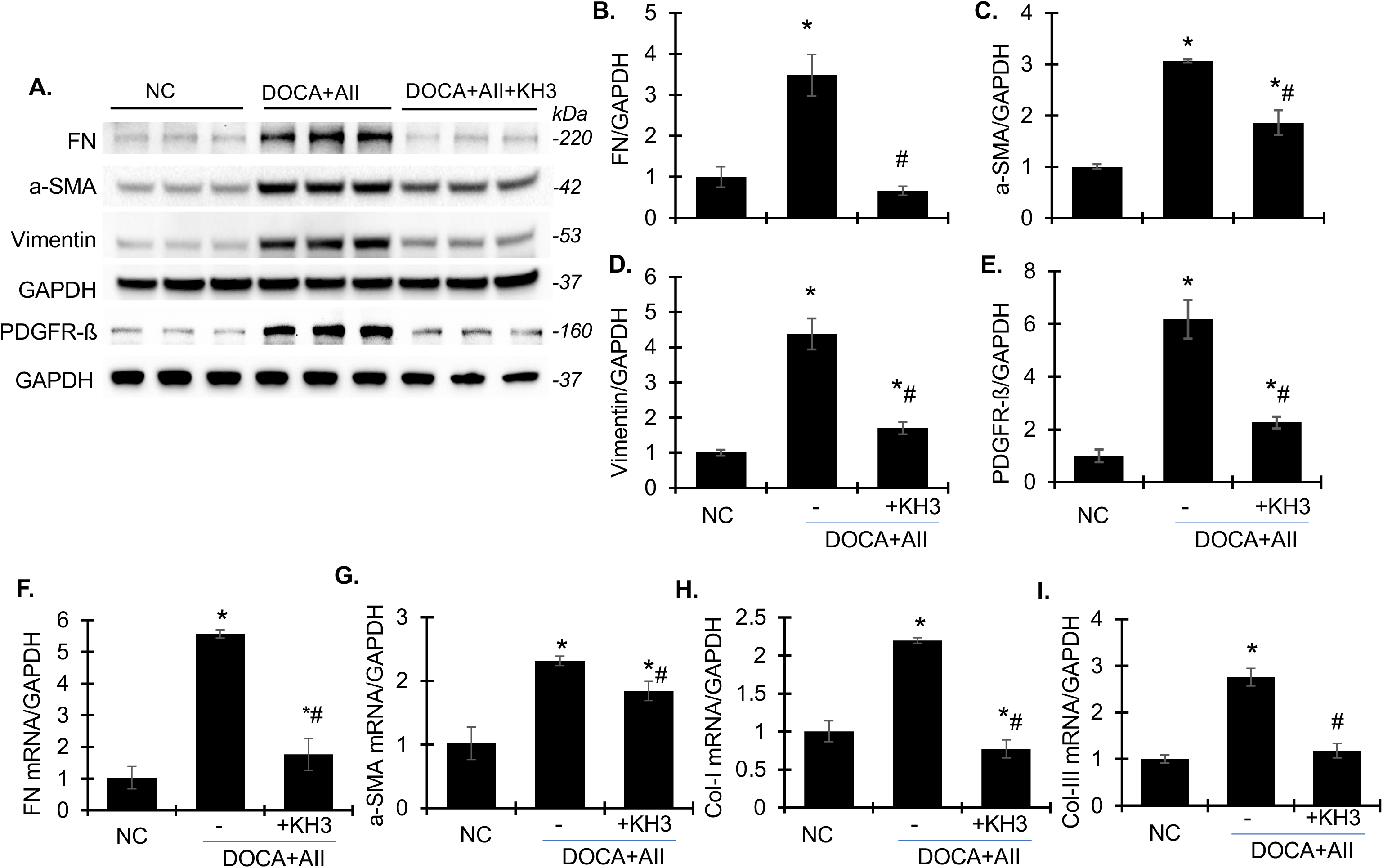
Treatment with KH3 reduces renal protein and mRNA expression of fibrotic markers in DOCA + Ang II-induced CKD mice. (**A**) Representative Western blots showing protein expression of FN, a-SMA, vimentin, PDGFR-ß, and GAPDH in kidneys from normal control, DOCA + Ang II treated, KH3-treated DOCA + Ang II mice. Molecular weight was indicated on the right. (**B-E)** Density quantification of FN (**B**), a-SMA (**C**), Vimentin (**D**), and PDGFR-ß (**E**) protein levels normalized to GAPDH. (**F-I)** Renal mRNA expression of FN (**F**), a-SMA (**G**), Col-I (**H**), and Col-III (**I**), as determined by quantitative RT-PCR. Data are presented as mean ± SD. *P<0.05 vs normal control (NC). #P<0.05, vs. DOCA+AII.

#### HuR inhibition preserves podocyte integrity

Normal control mice displayed strong linear nephrin (NP) and podocin (PD) staining along glomerular capillary loops (Fig. 5 A–B). DOCA + Ang II infusion reduced and disrupted nephrin and podocin staining (−27.2% and −22.6%, respectively; Fig. 5 C–D). KH3 largely restored nephrin and podocin expression toward control levels (Fig. 5 A–D), consistent with preserved slit diaphragm integrity and a mechanistic contribution to reduced albuminuria.

**Figure 5.**
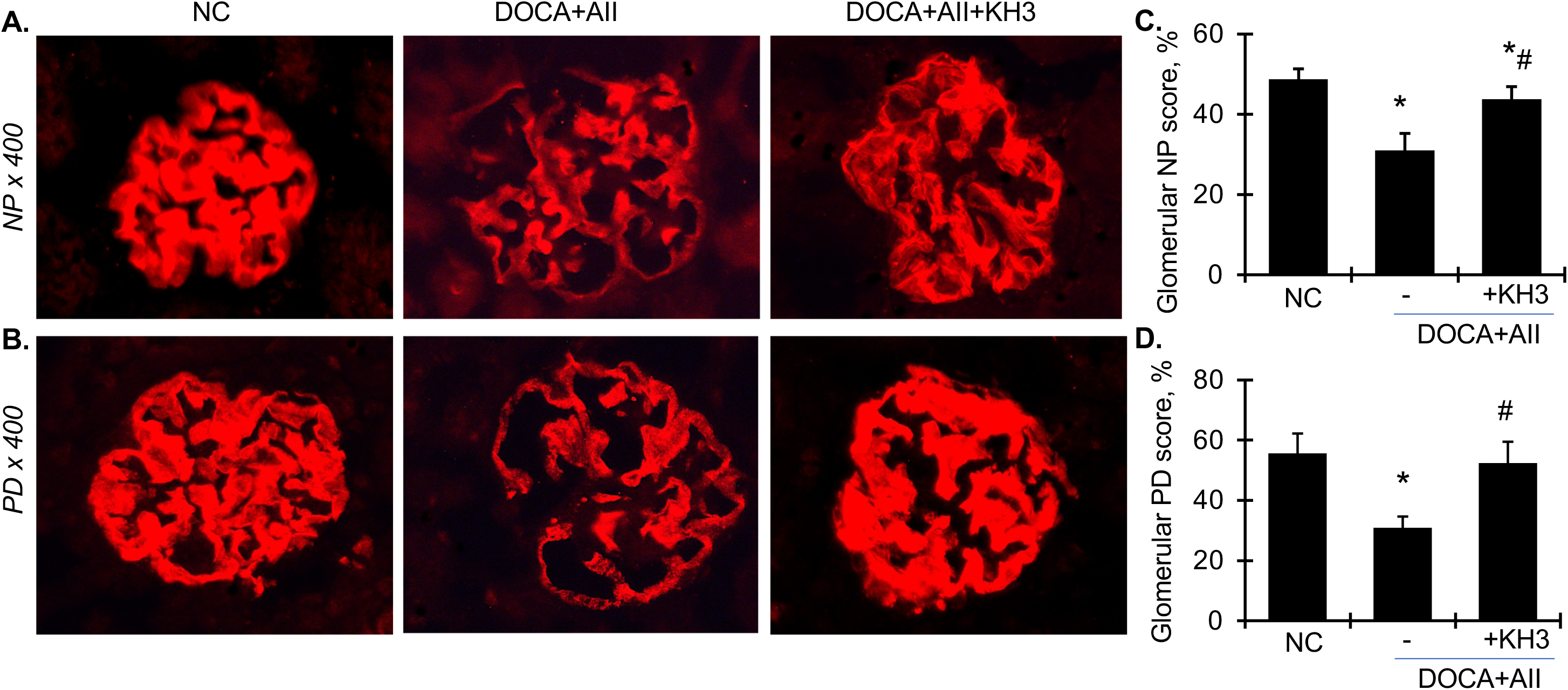
Treatment with KH3 attenuates glomerular podocyte injury in DOCA + Ang II-induced CKD mice. (**A-B**) Representative confocal images of kidney sections immunostained for nephrin (NP, red) (**A**) and podocin (PD, red) (**B**). Images were acquired at x400 magnification. (**C-D**) Quantitative analysis of NP (**C**) and PD (**D**) staining intensity performed using image-J. Data are presented as mean ± SD. *P<0.05 vs normal control (NC). #P<0.05, vs. DOCA+AII.

#### HuR inhibition reduces tubular injury

Because tubulointerstitial injury strongly predicts CKD outcomes ^24^, we assessed tubular injury markers. DOCA + Ang II infusion markedly increased renal Kim-1 and NGAL mRNA levels (Fig. 6 A–B). KH3 reduced Kim-1 and NGAL expression by ∼85% and ∼90%, respectively, although levels remained slightly above controls. Kim-1 and osteopontin (OPN) proteins were increased in diseased kidneys and were markedly reduced by KH3 toward control levels (Fig. 6 C–E). These data indicate that HuR inhibition substantially reverses tubular injury in hypertensive CKD.

**Figure 6.**
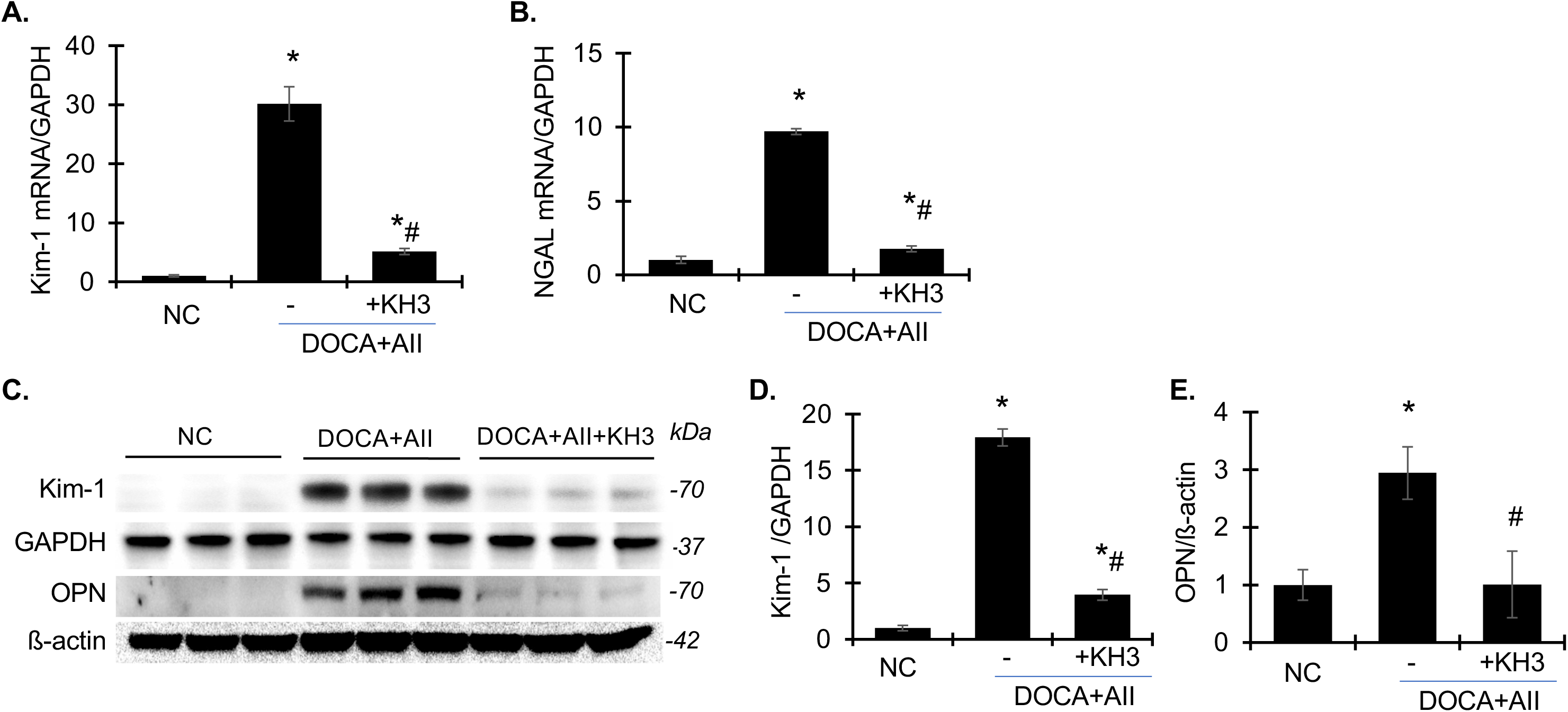
Treatment with KH3 reduces mRNA and protein expression of markers of tubular injury in DOCA + Ang II induced CKD mice. (**A-B**). Renal mRNA expression of Kim-1 (**A**) and NGAL (**B**), as determined by quantitative RT-PCR. (**C**) Representative Western blots showing protein expression of Kim-1, OPN, GAPDH, and ß-actin in kidneys from normal control, DOCA + Ang II treated, KH3-treated DOCA + Ang II mice. Molecular weight was indicated on the right. (**D-E**) Densitometric quantification of Kim-1 (**D**) and OPN (**E**) protein levels normalized to GAPDH and ß-actin, respectively. Data are presented as mean ± SD. *P<0.05 vs normal control (NC). #P<0.05, vs. DOCA+AII.

#### HuR inhibition suppresses inflammatory, oxidative stress, maladaptive repair, and profibrotic signaling

Western blotting showed increased NF-κBp65 in DOCA + Ang II-injured kidneys, which was suppressed by KH3 (Fig. 7 A–B). Nox2 protein was also increased and reduced to near baseline by KH3 (Fig. 7 A, C). AKT phosphorylation increased in DOCA + Ang II-injured kidneys and was reduced by ∼65% with KH3 (Fig. 7 A, D). Correspondingly, KH3 reduced renal mRNA expression of NF-κBp65, Nox2, MCP-1, and IL-6 (Fig. 7 E–H). F4/80 staining demonstrated increased macrophage infiltration in glomerular and interstitial compartments, which was markedly reduced by KH3 (Fig. 7 I–J). TGF-β1 and Wisp1 mRNA and protein were increased in diseased kidneys and attenuated by KH3 (Fig. 8 A–E). Collectively, KH3 suppresses NF-κB–driven inflammation, Nox2-dependent oxidative stress, AKT-mediated maladaptive repair, and TGF-β1/Wisp1 profibrotic signaling.

**Figure 7.**
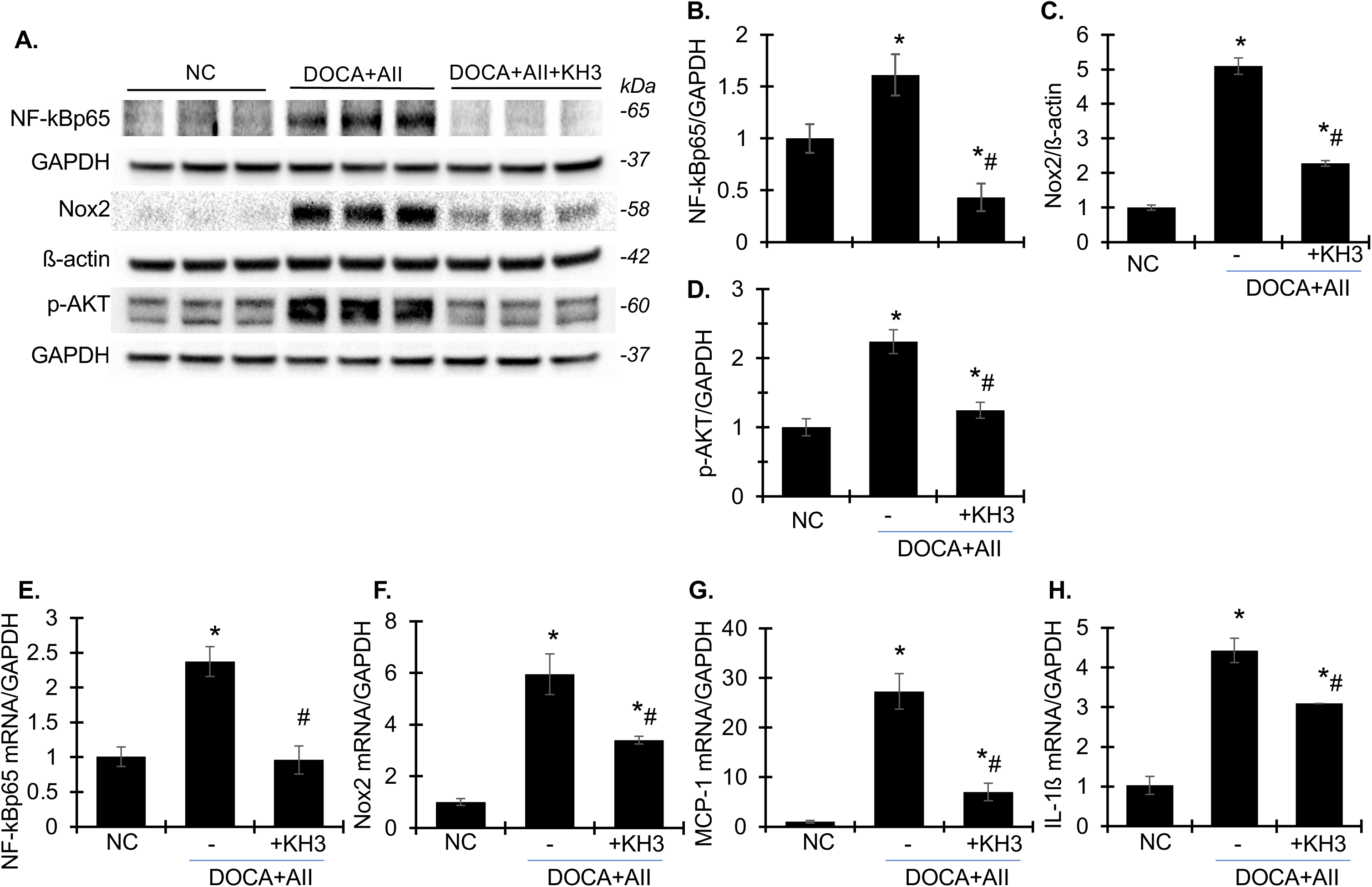
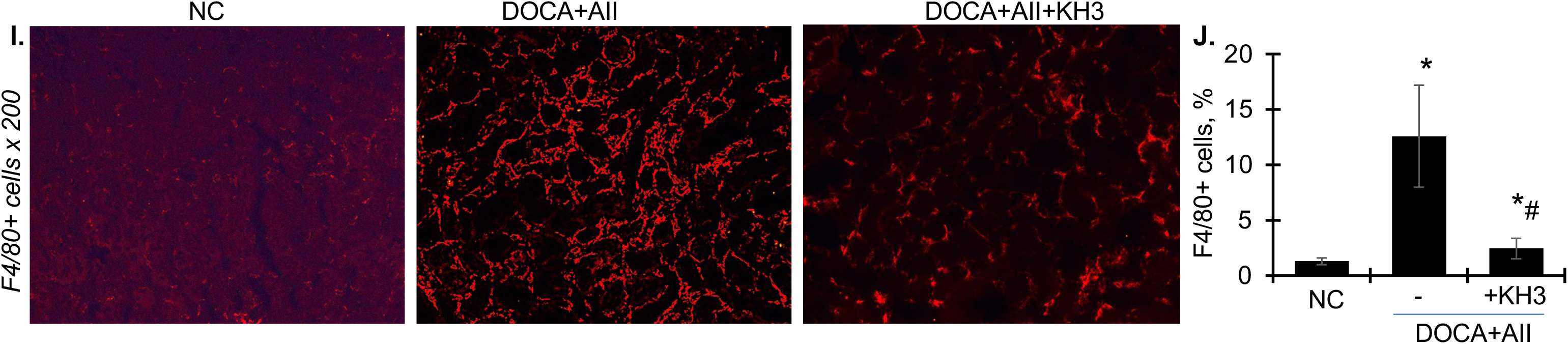
Treatment with KH3 suppresses renal pro-inflammatory, oxidative stress, and maladaptive signaling pathways and reduces macrophage infiltration in DOCA + Ang II-induced CKD mice. (**A**) Representative Western blots showing protein expression of NF-kBp65, Nox2, pAKT, ß-actin and GAPDH in kidneys from normal control, DOCA + Ang II treated, KH3-treated DOCA + Ang II mice. Molecular weight was indicated on the right. (**B-D**) Densitometric quantification of NF-kBp65 (**B**), Nox2 (**C**) and pAKT (**D**) protein levels normalized to GAPDH or ß-actin, as indicated. (E-H) Renal mRNA expression of NF-kBp65 (**E**), Nox2 (**F),** MCP-1 (**G**), and IL-1ß (**H**), as determined by quantitative real-time RT-PCR. Relative mRNA levels were calculated after normalization of GAPDH. (**I**) Representative confocal images of kidney sections immunostained for F4/80+macroopaghes (red). Images were acquired at x200 magnification. (**J**) Quantitative analysis of F4/80+ macrophage staining intensity performed using image-J. Data are presented as mean ± SD. *P<0.05 vs normal control (NC). #P<0.05, vs. DOCA+AII.

**Figure 8.**
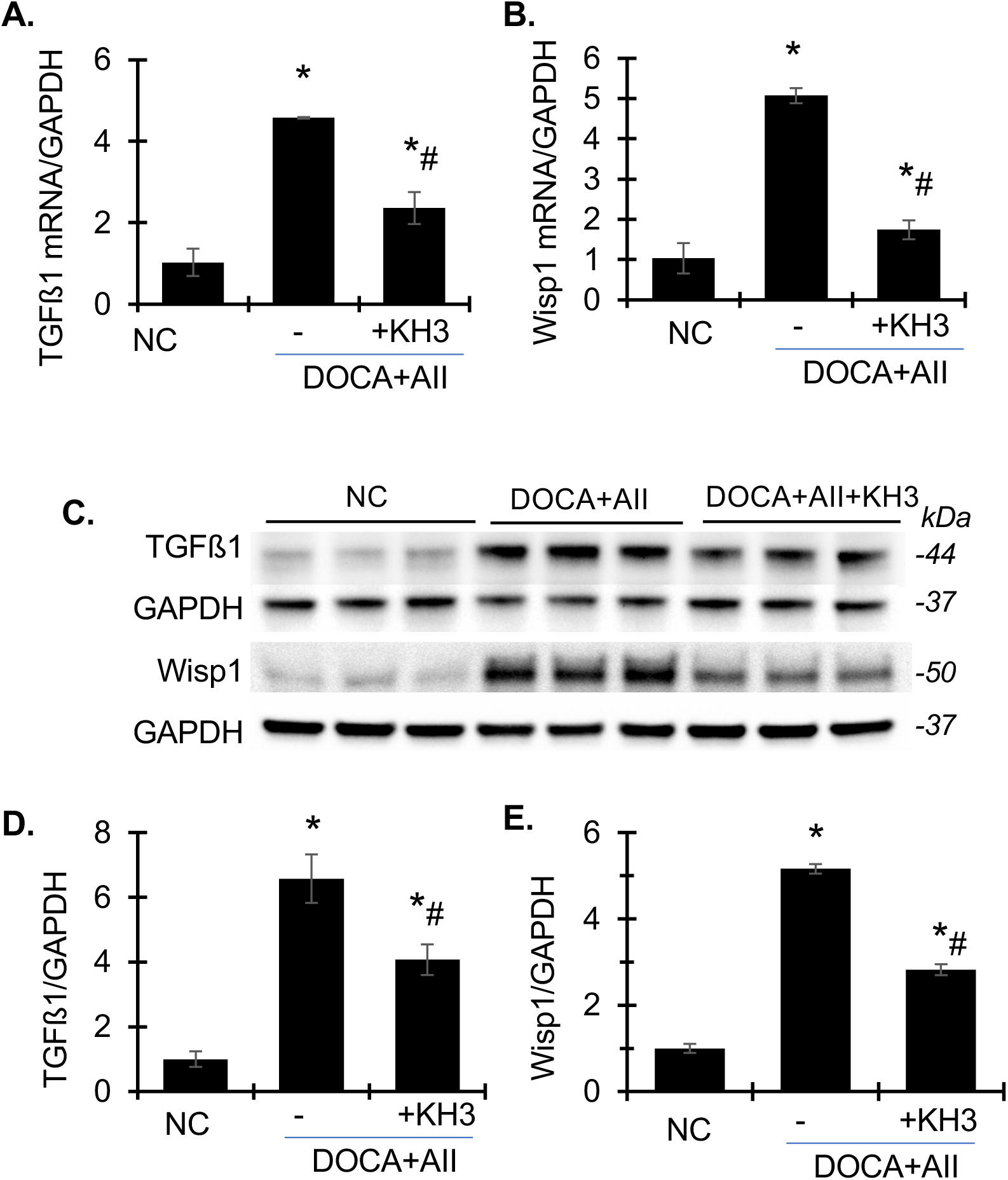
Treatment with KH3 attenuates renal TGFß1 and Wisp1 signaling in DOCA + Ang II-induced CKD mice. **(A-B)** Renal mRNA expression of TGFß1 (**A**) and Wisp1 (**B**), as determined by quantitative real-time RT-PCR. Relative mRNA levels were calculated after normalization of GAPDH. (**C**) Representative Western blots showing protein expression of TGFß1, Wisp1, and GAPDH in kidneys from normal control, DOCA + Ang II treated, KH3-treated DOCA + Ang II mice. Molecular weight was indicated on the right. (**D-E**) Densitometric quantification of TGFß1 (**D**) and Wisp1 (**E**) protein levels normalized to GAPDH. Data are presented as mean ± SD. *P<0.05 vs normal control (NC). #P<0.05, vs. DOCA+AII.

### HuR inhibition modulates core regulators of hypertension

Given the blood pressure reduction, we assessed renal sodium-handling proteins. First, SGLT2 protein increased in DOCA + Ang II-infused kidneys and was reduced by ∼32% with KH3; GLP-1R was unchanged among groups (Fig. 9 A–C). In DOCA + Ang II-infused kidneys, SGLT2 staining increased not only in proximal tubules but also in renal arterial vessels, and both compartments were reduced by KH3 (Fig. 9I). Second, in this model, ENaC subunits showed a non-uniform pattern (ENaC-β unchanged, ENaC-α modestly reduced, and ENaC-γ markedly increased) (Fig. 9 D–H), consistent with disease-associated distal nephron remodeling rather than a coordinated physiologic mineralocorticoid transcriptional response. Notably, KH3 did not reduce the pronounced ENaC-γ induction, whereas it suppressed SGLT2 expression and partially normalized ENaC-α, suggesting that HuR inhibition influences sodium-handling predominantly through SGLT2-associated pathways and broader maladaptive inflammatory/profibrotic signaling rather than global suppression of ENaC (Fig. 9 A–H). These findings also prompted further investigation of SGLT2.

**Figure 9.**
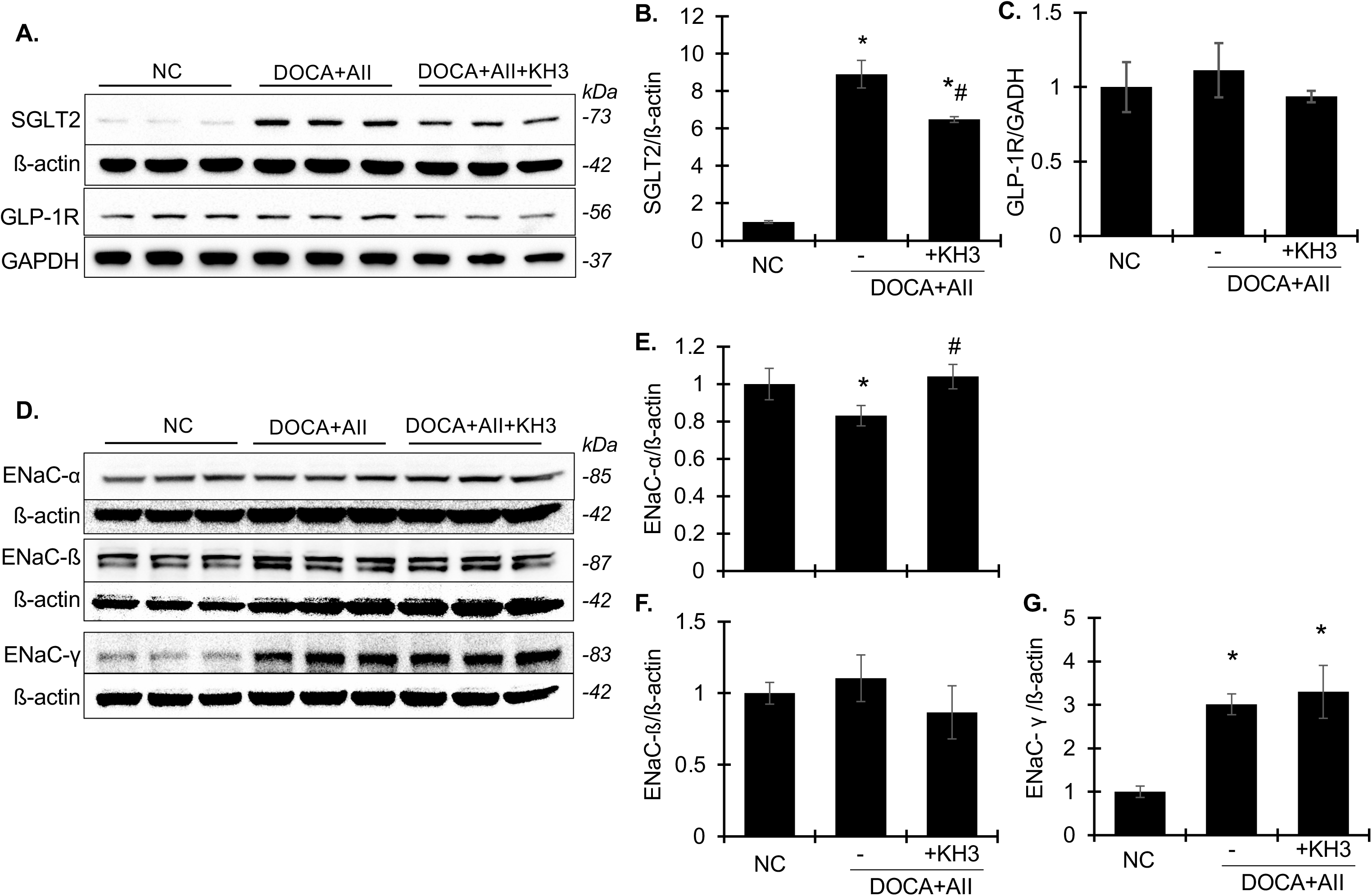
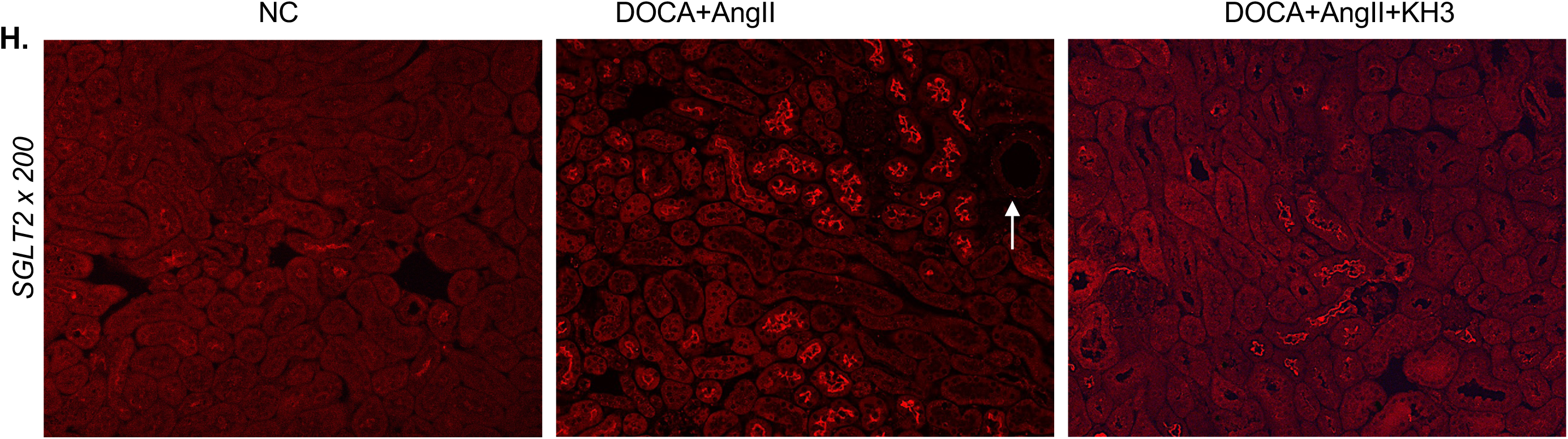
KH3 treatment modulates renal expression of SGLT2, GLP-1R, and ENaC subunits in DOCA + Ang II-induced CKD mice. (**A, D**) Representative Western blots showing renal protein expression of SGLT2 and GLP-1R (**A**), as well as ENaC subunits a, ß, and γ (**D**), in kidneys from normal control, DOCA + Ang II treated, KH3-treated DOCA + Ang II mice. β-actin or GAPDH served as loading controls, as indicated. Molecular weight was indicated on the right. (**B-C, E-G**) Densitometric quantification of SGLT2 (**B**), GLP-1R (**C**), ENaC-a (**E**), ENaC-ß (**F**), and ENaC-γ (**G**) protein levels normalized to ß-actin or GAPDH, as indicated. Data are presented as mean ± SD. (**H**) Representative confocal images of kidney sections immunostained for SGLT2 (red). Images were acquired at x200 magnification. Arrows indicate vascular SGLT2 staining. Data are presented as mean ± SD. *P<0.05 vs normal control (NC). #P<0.05, vs. DOCA+AII.

### SGLT2 regulation in arteries and VSMCs

In aortas, SGLT2 mRNA was detectable and increased in DOCA + Ang II-infused mice, along with upregulation of AT1 receptor (AT1R), TGF-β1, Wisp1, NF-κBp65, ID3, and prorenin receptor (PRR). KH3 normalized the expression of all transcripts except PRR (Fig. 10 A–H). These data demonstrate that DOCA + Ang II promotes a proinflammatory and profibrotic program in the arterial wall, which is largely dependent on HuR activity.

**Figure 10.**
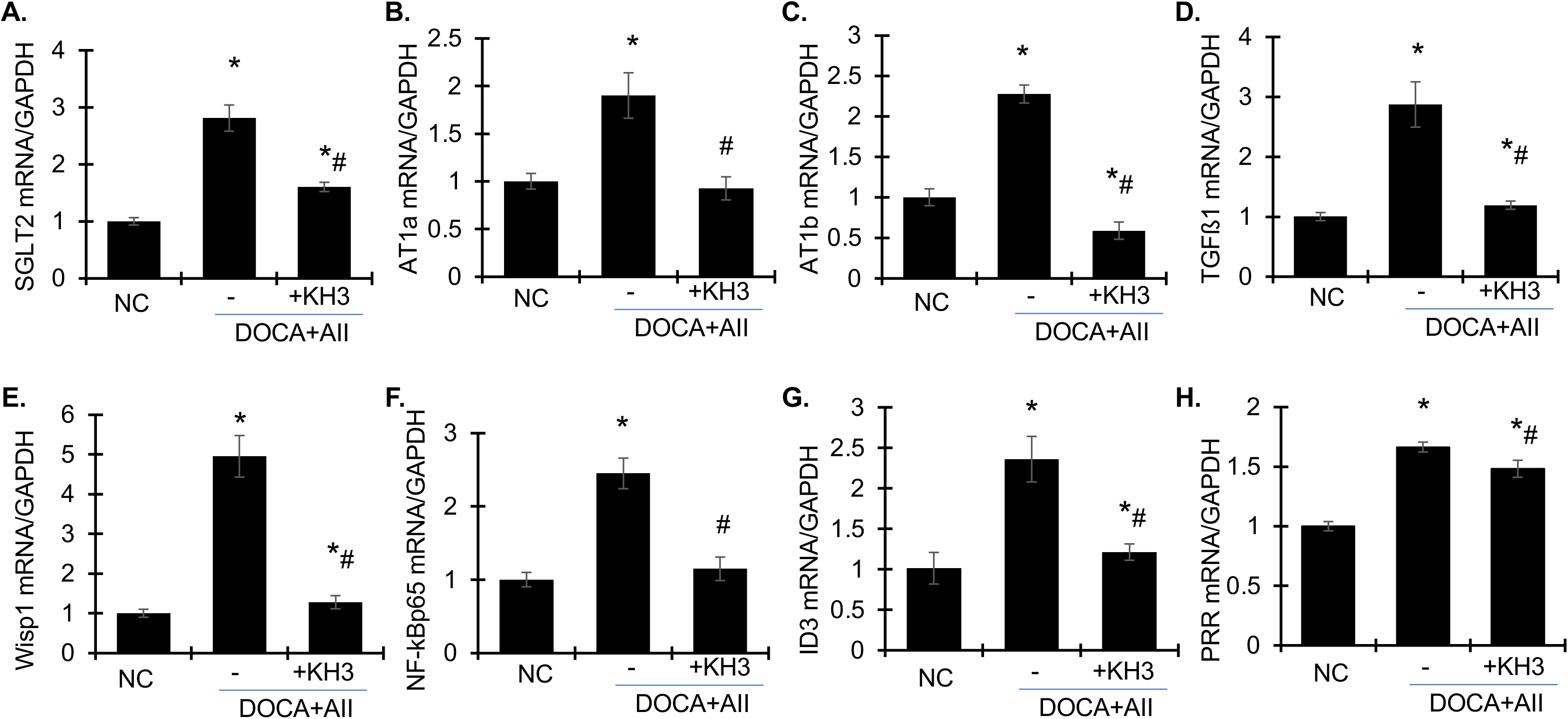
KH3 treatment attenuates artery mRNA expression of SGLT2, AT1 receptor, and profibrotic / proinflammatory markers in DOCA + Ang II-induced CKD mice. **(A-H)** Artery mRNA expression of SGLT2 (**A**), AT1a (**B),** AT1b (**C**), TGFß1 (**D**), Wisp1 (**E**), NF-kBp65 (**F**), ID3 (**G**), and prorenin receptor (PRR) (**H**), as determined by quantitative real-time RT-PCR. Relative mRNA levels were calculated after normalization of GAPDH. Data are presented as mean ± SD. *P<0.05 vs normal control (NC). #P<0.05, vs. DOCA+AII.

Primary mVSMCs expressed SGLT2 at lower levels than mPTCs by RT-qPCR and Western blotting (Fig. 11 A–B). Immunofluorescence suggested perinuclear / nuclear-associated staining (Fig. 11C). Ang II and TGF-β1 increased SGLT2 mRNA in mVSMCs along with Wisp1 and IL-6 (Fig. 11 D–I); TGF-β1 was then selected for mechanistic studies. TGF-β1 increased cytoplasmic HuR and/or HuR translocation, which was reversed by KH3 (Fig. 12 A-C). KH3 strongly suppressed TGF-β1–induced SGLT2 protein expression, reducing levels below unstimulated controls, supporting HuR-dependent regulation of SGLT2 under profibrotic conditions (Fig. 13 A-B). TGF-β1 also increased NF-κBp65, Wisp1, and α-SMA; these effects were mitigated by SGLT2 inhibitor dapagliflozin (Fig. 13 C-G). Together, these results support a HuR-dependent SGLT2 program that contributes to inflammatory and profibrotic VSMC activation.

**Figure 11.**
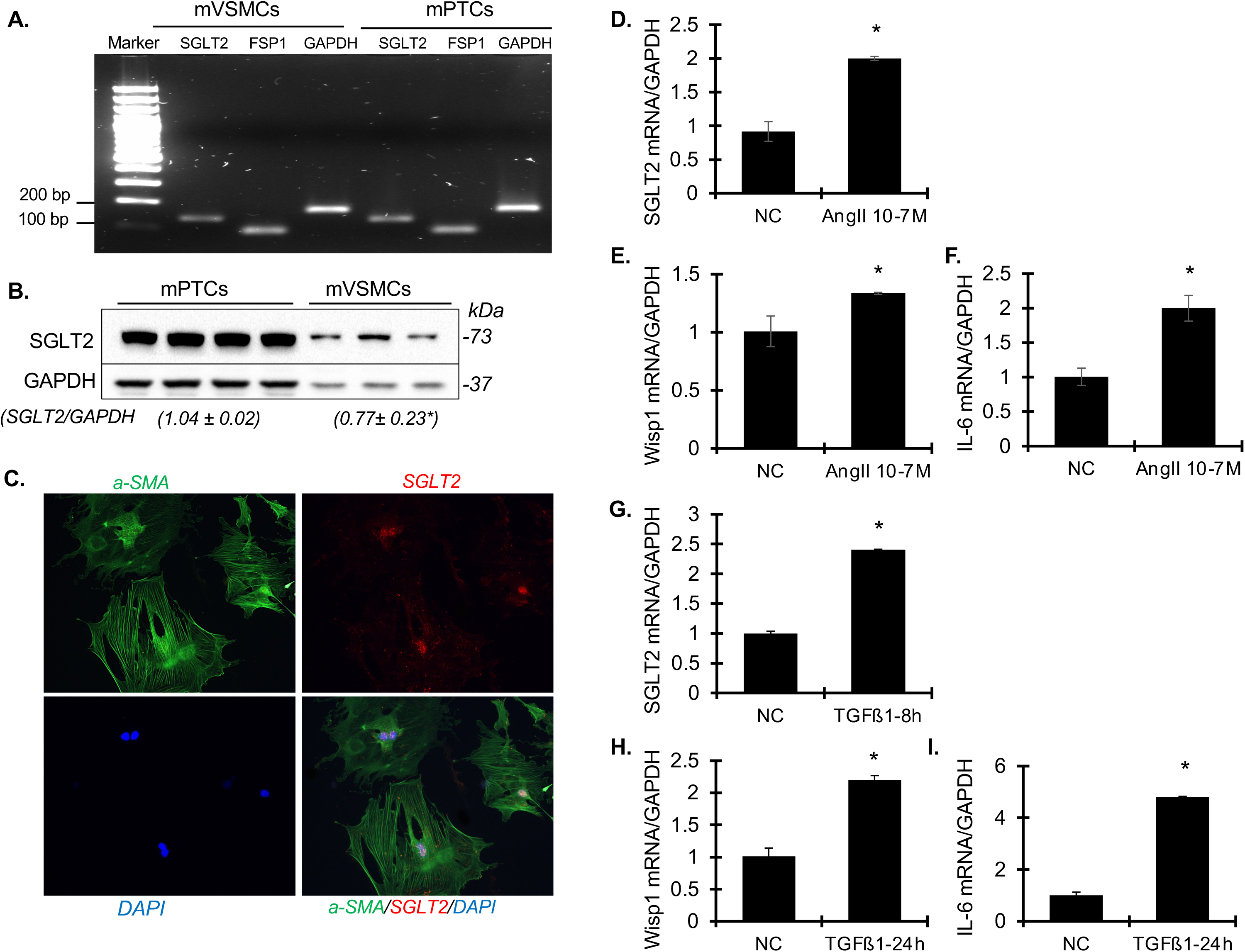
Vascular smooth muscle cells (VSMCs) express SGLT2 and its expression is regulated by angiotensin II (Ang II) and TGFß1. (**A**) SGLT2 mRNA in primary cultured mouse VSMCs compared with primary cultured mouse PTCs, with FSP1 and GAPDH shown as controls. mRNA expression was determined by real time RT-PCR and visualized on a 2% agarose gel. DAN ladders (Mr) are shown. (**B**) Representative Western blots showing SGLT2 protein levels in cultured mPTCs and mVSMCs. Densitometric quantification of SGLT2 protein levels normalized to GAPDH is shown below the blots. (**C**) Representative immunofluorescence images of cultured mVSMCs stained for a-SMA (green), SGLT2 (red), and nuclei (DAPI, blue). Images were acquired at x400 magnification. (**D-I**) mRNA expression of SGLT2 (**D, G**), Wisp1 (**E, H),** and IL-6 (**F, I**) in mVSMCs following stimulation with Ang II (**D-F**) or TGFß1 (**G-I**), as determined by quantitative real-time RT-PCR. Relative mRNA levels were calculated after normalization of GAPDH. Data are presented as mean ± SD. *P<0.05 vs untreated control cells (normal control, NC).

**Figure 12.**
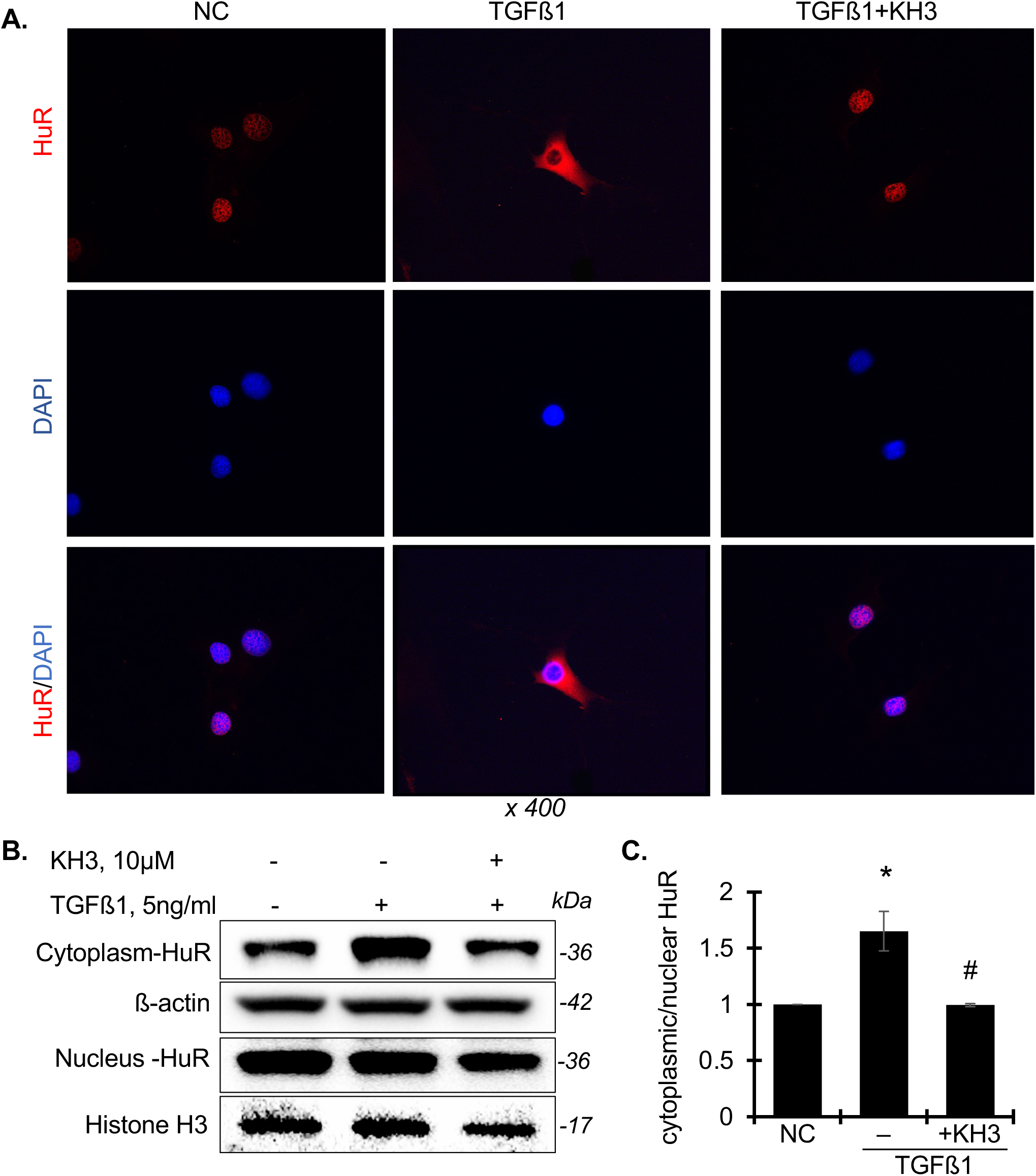
KH3 treatment abrogates TGFß1-induced cytoplasmic location and expression of HuR in primary cultured mouse vascular smooth muscle cells (mVSMCs). (**A**) Representative immunofluorescent staining images showing HuR (red) and nuclei (DAPI, blue) in the un-treated mVSMCs (normal control, NC), mVSMCs treated with TGFß1 alone or mVSMCs treated with TGFß1 in combination with KH3. Arrows indicate increased cytoplasmic HuR staining. (**B**) Representative Western blots showing HuR protein levels in cytoplasmic and nuclear fractions from untreated and treated mVSMCs. β-actin and histone proteins were used as loading controls for cytoplasmic and nuclear fractions, respectively. (**C**) Densitometric quantification of the ratio of cytoplasmic to nuclear HuR protein expression. Values are expressed relative to untreated cells, which was set to unity. Data are presented as mean ± SD. *P<0.05, vs. untreated control (NC); #P<0.05, vs. TGFß1 alone-treated cells.

**Figure 13.**
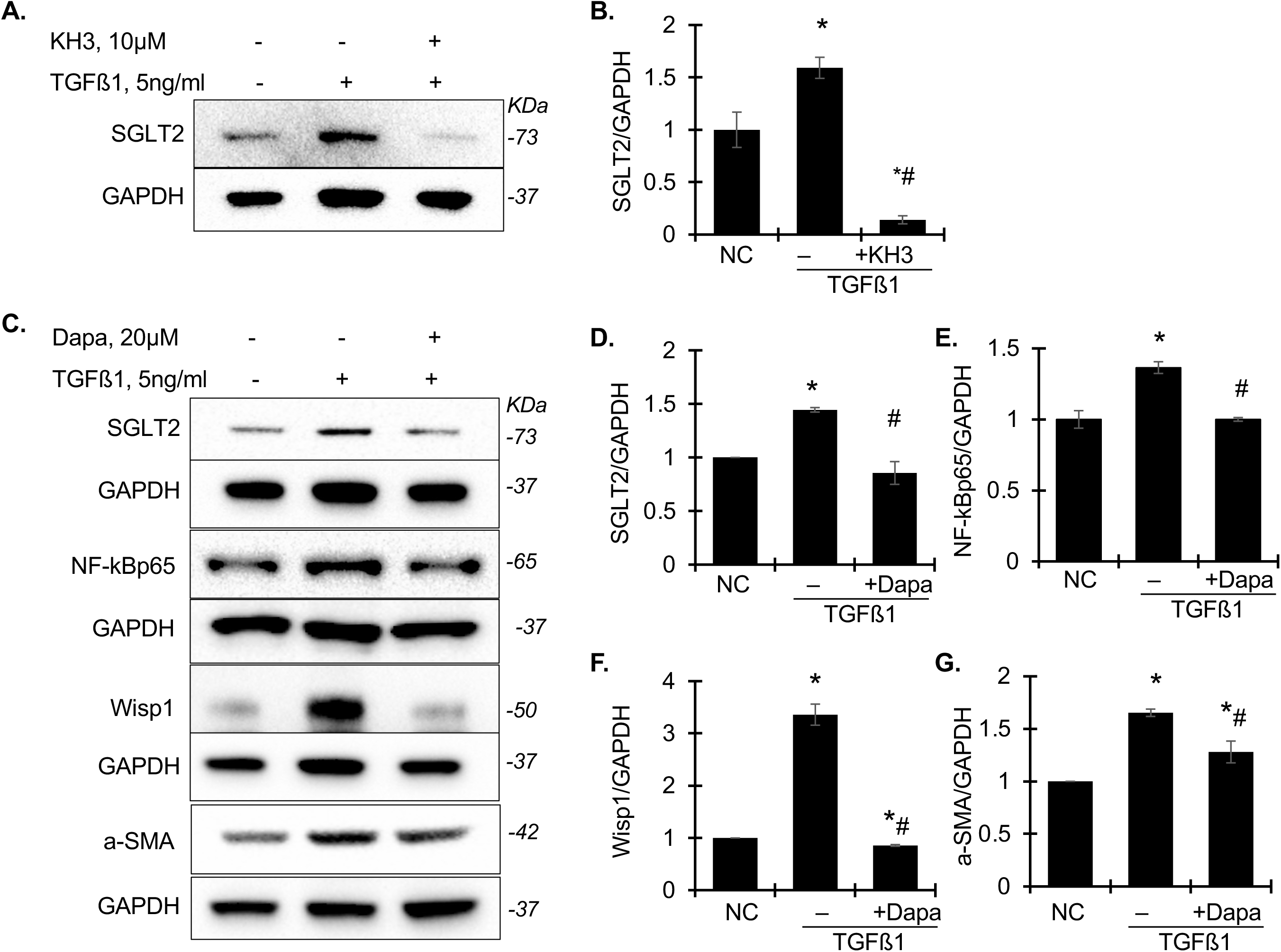
HuR inhibition abrogates TGFß1-induced SGLT2 expression and downstream proinflammatory and profibrotic signaling in cultured mVSMCs. (**A**) Representative Western blots showing SGLT2 and GAPDH protein expression in cultured mVSMCs following TGFß1 stimulation in absence or presence of the HuR inhibitor KH3. (**B**) Densitometric quantification of SGLT2 protein levels from (A), normalized to GAPDH and expressed relative to untreated control cells (set to unity). (**C**) Representative Western blots showing protein expression of SGLT2, NF-kBp65, Wisp1, a-SMA and GAPDH protein expression in cultured mVSMCs following TGFß1 stimulation in absence or presence of SGLT2 inhibitor dapagliflozin (Dapa). (**D-G**) Densitometric quantification of SGLT2 (**D**), NF-kBp65 (**E**), Wisp1 (**F**), and a-SMA (**G**), protein levels from (C), normalized to GAPDH and expressed relative to untreated control cells. Data are presented as mean ± SD. *P<0.05, vs. untreated control (NC); #P<0.05, vs. TGFß1 alone-treated cells.

### HuR and SGLT2 inhibition reduce Ang II–mediated vasoconstriction

Mesenteric artery characteristics i.e., initial width, width at Lmax, length, and tension development at Lmax were similar across groups (Supplemental Table S3). Likewise, maximal responses to non-receptor mediated vasocontraction via KCl and a-1 receptor -mediated vasocontraction via PE, were not different among groups. In contrast, Ang II-induced AT1R-mediated vasocontraction in the presence of vehicle was blunted by KH3, dapagliflozin (Dapa), and to a greater extent by their combination (Fig. 14 A-B). Endothelium-dependent relaxation to acetylcholine and endothelium-independent relaxation to sodium nitroprusside were similar across groups, indicating preserved endothelial and vascular smooth muscle integrity.

**Figure 14.**
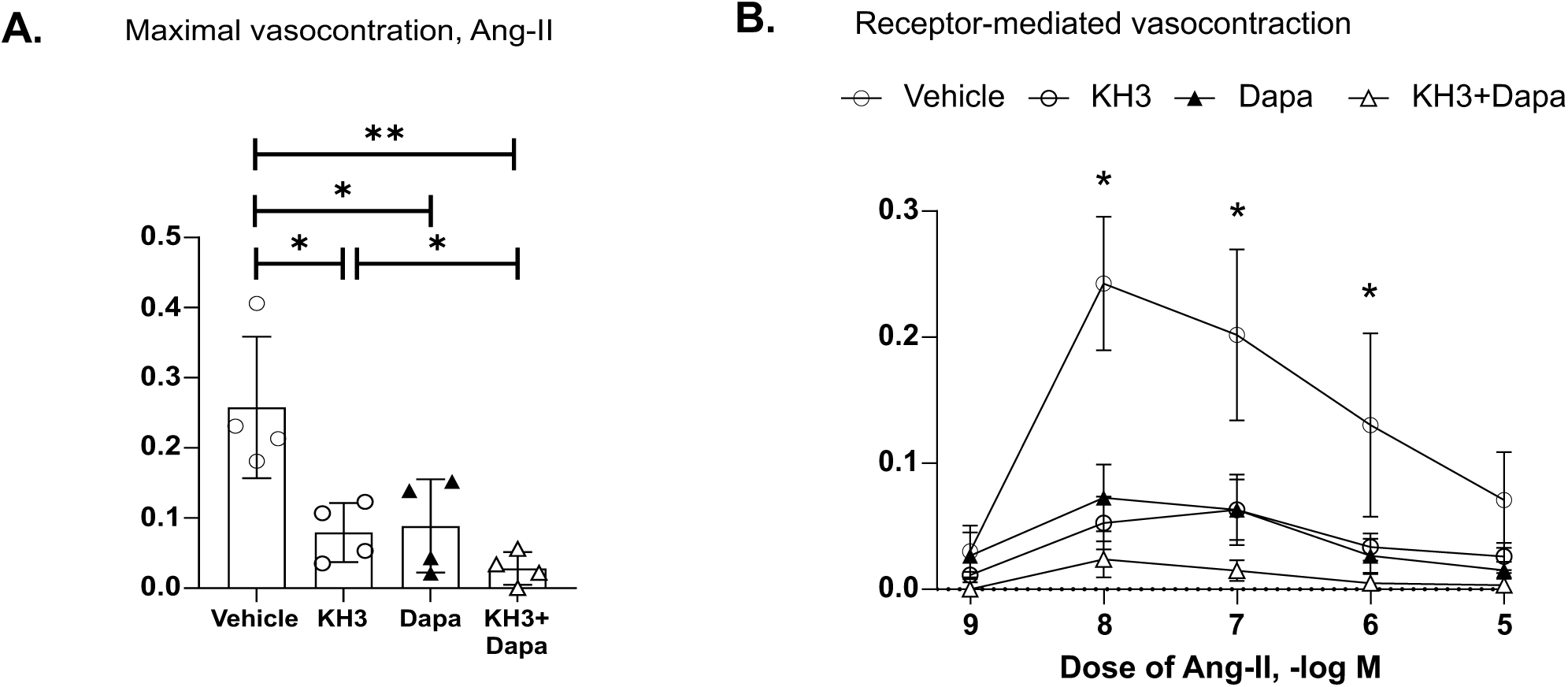
HuR and SGLT2 inhibition independently and additively attenuate AT1 receptor-mediated vasocontraction in mesenteric resistance arteries. (**A**) Concentration-response curves showing angiotensin II (Ang II)-induced vasoconstriction of mesenteric resistance arteries following 120-min incubation with vehicle (V; PBS), the HuR inhibitor KH3 (10 μM), the SGLT2 inhibitor dapagliflozin (Dapa; 20 μM), or the combination of KH3 and Dapa. Vasoconstrictor responses to cumulative Ang II concentrations were significantly attenuated by KH3, Dapa, and their combination compared with vehicle treatment. Each data point represents one artery segment per mouse from four mice per group. Data are presented as mean ± SEM. *P < 0.05, **P < 0.01 compared with indicated control groups. (**B**) Maximal Ang II–induced vasoconstriction derived from the concentration–response curves in (A). Maximal contractile responses were similarly blunted by KH3, Dapa, and their combination compared with vehicle treatment. Individual data points (circles) represent single artery segments from individual mice. *P < 0.05 vs. vehicle (V).

## DISCUSSION

In the present study, we identify HuR as a previously underappreciated integrative regulator of hypertensive CKD, linking renal inflammation and fibrosis with vascular dysfunction and blood pressure control. Using the DOCA + Ang II model, we demonstrate robust HuR upregulation at systemic and renal levels, including in glomerular, tubular, vascular, and infiltrating inflammatory cells. Pharmacologic inhibition of HuR with KH3 not only attenuated renal inflammation, fibrosis, podocyte injury, and tubular damage, but also significantly lowered blood pressure-an effect not previously attributed to HuR inhibition. These findings position HuR as a central upstream mediator of kidney-vascular crosstalk in hypertensive CKD.

Our data extend prior work implicating HuR in renal inflammation and fibrosis by demonstrating broad pathogenic roles across multiple renal compartments in hypertensive injury. KH3 consistently suppressed proinflammatory pathways (NF-κB, MCP-1, IL-6), oxidative stress signaling (Nox2), and profibrotic mediators (TGF-β1, Wisp1), and reduced AKT phosphorylation associated with maladaptive repair. Importantly, HuR inhibition preserved podocyte slit diaphragm integrity and markedly reduced tubular injury markers (Kim-1, NGAL, OPN), supporting HuR as a key post-transcriptional regulator of disease-sustaining gene networks in CKD.

A major and unexpected finding is that HuR inhibition significantly attenuated hypertension in DOCA + Ang II-treated mice. To our knowledge, this is the first demonstration that pharmacologic HuR inhibition lowers blood pressure in vivo. This antihypertensive effect was accompanied by reduced polydipsia, polyuria, and albuminuria, supporting a true improvement in systemic hemodynamics rather than a secondary consequence of renal protection alone.

Mechanistically, HuR inhibition selectively modulated renal sodium-handling pathways, most notably through suppression of SGLT2 expression in renal tubular cells and vessel cells, with minimal effects on regulation of ENaC subunits. In parallel, KH3 treatment blunted AT1R-mediated vasocontraction in mesenteric resistance arteries to an extent comparable to SGLT2 inhibition, responses that were exacerbated by their combination. These findings indicate that HuR may influence blood pressure through both renal and extrarenal mechanisms.

Notably, we provide evidence that SGLT2 is expressed not only in renal tubular epithelial cells but also in VSMCs, where expression is upregulated in hypertensive conditions. HuR inhibition robustly suppressed SGLT2 in arteries and cultured VSMCs, together with Ang II responsive and profibrotic genes including AT1R, TGF-β1, Wisp1, NF-κB, and ID3, supporting a previously unrecognized role for SGLT2 in vascular stress signaling. In vitro, TGF-β1 induced HuR cytoplasmic accumulation and SGLT2 upregulation in VSMCs, both reversed by KH3. Direct SGLT2 inhibition with dapagliflozin similarly suppressed inflammatory and contractile markers. Ex vivo, both KH3 and dapagliflozin attenuated Ang II-induced vasocontraction, with additive effects when combined. Together, these data identify a HuR-SGLT2-VSMC axis that contributes to vascular hypercontractility and hypertension in CKD.

At first glance, these results may contrast with reports that constitutive VSMC-specific HuR deletion causes hypertension ^25^. Several distinctions likely explain this discrepancy. Prior studies employed non-inducible, cell-specific HuR deletion, which may disrupt vascular development, smooth muscle differentiation, and baseline vessel architecture. Because HuR regulates transcripts involved in cytoskeletal organization and contractile protein expression ^25^, its lifelong absence may predispose to developmental vascular abnormalities and compensatory hypertension. In contrast, our study used pharmacologic inhibition in adult mice with established disease, targeting disease-associated HuR overactivation rather than basal HuR function. Moreover, KH3 did not abolish HuR expression but normalized its disease-induced upregulation, potentially preserving physiological HuR functions while suppressing maladaptive signaling. In addition, HuR inhibition acted in a multicellular context, simultaneously modulating renal tubular injury, vascular inflammation, macrophage infiltration, and VSMC contractility-effects not captured by single cell–type knockout models.

SGLT2 has traditionally been viewed as a proximal tubular transporter. However, accumulating evidence supports context-dependent SGLT2 expression in extra-renal and vascular compartments. Recent studies report increased SGLT2 expression in arterial endothelium, vascular smooth muscle, coronary microvasculature, and cardiomyocytes under inflammatory conditions ^26^. Proinflammatory cytokines (IL-1β, IL-6, TNF-α) can increase SGLT2 expression in cultured endothelial cells and promote NF-κB activation and oxidative stress, whereas SGLT2 inhibition or renin angiotensin aldosterone system (RAAS) blockade reduces ROS and improves redox balance ^26^. These observations provide a plausible framework for some of the rapid cardiovascular benefits observed clinically with SGLT2 inhibitors.

Consistent with this paradigm, experimental studies suggest that SGLT2 expression in VSMCs may be induced by pathological stimuli and contribute to vascular remodeling ^27^. Dapagliflozin has been reported to attenuate vascular calcification and osteogenic reprogramming in VSMCs, supporting a direct vascular component that extends beyond renal hemodynamics ^28^. Our findings align with these reports and further position SGLT2 as part of a broader stress-response module in diseased vasculature that is regulated upstream by HuR.

At the same time, the literature remains divided on whether cardiovascular benefits of SGLT2 inhibitors require direct SGLT2 blockade in non-renal cells. Several mechanistic studies implicate alternative sodium-handling targets (e.g., NHE1, SMIT, SMVT) or systemic mechanisms such as natriuresis/ tubuloglomerular feedback, reduced sympathetic/ RAAS tone, metabolic reprogramming, and immunomodulation ^29–32^. Notably, SGLT2 inhibitors can reduce macrophage activation and recruitment, which may rapidly improve vascular inflammation and tone in CKD ^33^. In this context, our data suggest that HuR serves as an upstream integrator of inflammatory/profibrotic programs and a regulator of vascular SGLT2 induction, providing a mechanistic bridge between HuR inhibition and vascular functional improvement.

Collectively, these findings support a model in which extra-proximal tubule expression of SGLT2 is inducible in inflammatory and hypertensive states and may amplify RAAS-NF-κB-NADPH oxidase signaling in vascular cells, while SGLT2 inhibitors may confer both SGLT2-dependent and -independent vasculoprotective actions. Our study extends this framework by identifying HuR as an upstream post-transcriptional regulator of the vascular SGLT2 program and by demonstrating that HuR and SGLT2 inhibition converge to blunt Ang II driven vasoconstriction, with additive effects when combined. These findings may help explain clinically observed blood pressure and vascular benefits of SGLT2 inhibitors, including in non-diabetic CKD populations ^34–37^. Nonetheless, inducible VSMC- or endothelial cell-specific HuR and SGLT2 knockout models in hypertensive disease settings will be required to definitively delineate cell-autonomous versus non–cell-autonomous contributions to blood pressure regulation.

### Conclusions

We identify HuR as a central pathogenic regulator in hypertensive CKD that integrates renal injury, vascular dysfunction, and blood pressure control. We uncover a novel HuR-SGLT2 signaling axis in VSMCs and demonstrate that pharmacologic HuR inhibition lowers blood pressure while broadly suppressing inflammatory and fibrotic signaling. These findings reconcile apparent discrepancies with prior genetic studies and establish HuR inhibition as a promising mechanism-based therapeutic strategy for hypertensive CKD. The convergence of HuR and SGLT2 signaling in kidney and vasculature further suggests that HuR inhibition may complement SGLT2 inhibitors by simultaneously targeting multiple pathogenic pathways currently addressed by distinct drug classes.

### Perspectives

HuR has emerged as a central post-transcriptional regulator linking renal inflammation, vascular dysfunction, and blood pressure control in hypertensive CKD. By uncovering a HuR–SGLT2 signaling axis in vascular smooth muscle cells and demonstrating that HuR inhibition attenuates Ang II–mediated vasoconstriction, this study expands current paradigms of blood pressure regulation beyond classical renal sodium handling to include RNA-binding protein–mediated vascular stress signaling.

From a translational perspective, HuR represents an attractive upstream therapeutic target capable of simultaneously modulating multiple pathogenic pathways, including inflammation, oxidative stress, fibrosis, and vascular hypercontractility. These findings further suggest that HuR inhibition may complement existing antihypertensive strategies, including RAAS blockade and SGLT2 inhibition, and may help explain the blood pressure–lowering and vascular benefits of SGLT2 inhibitors observed even in non-diabetic CKD populations.

Future studies using inducible, cell-specific genetic models will be required to delineate the relative contributions of renal versus vascular HuR signaling to blood pressure regulation. Defining the temporal and spatial dynamics of HuR activation in hypertension may also facilitate the development of targeted combination therapies aimed at improving blood pressure control and reducing cardiovascular risk in CKD.

## AUTHOR CONTRIBUTIONS

YH created the study concept and design and supervised the whole study. LZ, ZW and YH performed animal and cell culture experiments and analyzed the data. ZF participated the exosomal and ENaC analyses. LZ wrote the first draft of manuscript. XW, JA, and LX designed, synthesized, purified and tested the KH3 in cancer cells and tumor growth. JDS and SM performed the vascular reactivity study and analyzed the related data. YH wrote and edited the manuscript and provided final approval and has overall responsibility for the published work. All authors read and approved the manuscript.

## SUPPLEMENTARY MATERIAL

Supplementary Tables S1 to S3.

Supplementary Methods

Unedited Western blots and gel images

